# Deep neural networks for predicting single cell responses and probability landscapes

**DOI:** 10.1101/2023.06.24.546373

**Authors:** Heidi E. Klumpe, Jean-Baptiste Lugagne, Ahmad S. Khalil, Mary J. Dunlop

## Abstract

Engineering biology relies on the accurate prediction of cell responses. However, making these predictions is challenging for a variety of reasons, including the stochasticity of biochemical reactions, variability between cells, and incomplete information about underlying biological processes. Machine learning methods, which can model diverse input-output relationships without requiring *a priori* mechanistic knowledge, are an ideal tool for this task. For example, such approaches can be used to predict gene expression dynamics given time-series data of past expression history. To explore this application, we computationally simulated single cell responses, incorporating different sources of noise and alternative genetic circuit designs. We showed that deep neural networks trained on these simulated data were able to correctly infer the underlying dynamics of a cell response even in the presence of measurement noise and stochasticity in the biochemical reactions. The training set size and the amount of past data provided as inputs both affected prediction quality, with cascaded genetic circuits that introduce delays requiring more past data. We also tested prediction performance on a bistable auto-activation circuit, finding that our initial method for predicting a single trajectory was fundamentally ill-suited for multimodal dynamics. To address this, we updated the network architecture to predict the entire distribution of future states, showing it could accurately predict bimodal expression distributions. Overall, these methods can be readily applied to the diverse prediction tasks necessary to predict and control a variety of biological circuits, a key aspect of many synthetic biology applications.

## Introduction

Mathematical models are crucial tools for predicting the responses of genetic circuits to signals and perturbations from the environment and for reliably engineering biological systems^1,2^. Models that can predict the effect of a gene knockout or overexpression, the response of a cell to a drug or change in carbon source, or the impact of an optogenetic input sequence can allow the design of more ambitious genetic circuits^3–13^. Thus, the ability to predict biological time-series data is critical to designing robust genetic circuits and can also provide novel insight into the behaviors of natural systems.

However, the nature of biological processes makes dynamic cell responses hard to predict (Figure 1A). The absolute number of many molecules in a cell is small, leading to stochastic behavior^14,15^. Processes such as gene expression are inherently bursty^16^, generating large fluctuations and keeping the cell far from steady-state. Moreover, the sharing of key resources, such as ribosomes^17^, across many different processes, as well as the pleiotropic and interconnected effects of many genes^18^, makes it difficult to fully explain the behavior of one system without accounting for other parts of the cell. Beyond these biological constraints, many technical challenges make such predictions difficult due to noisy and incomplete measurements of cell states.

**Figure 1:**
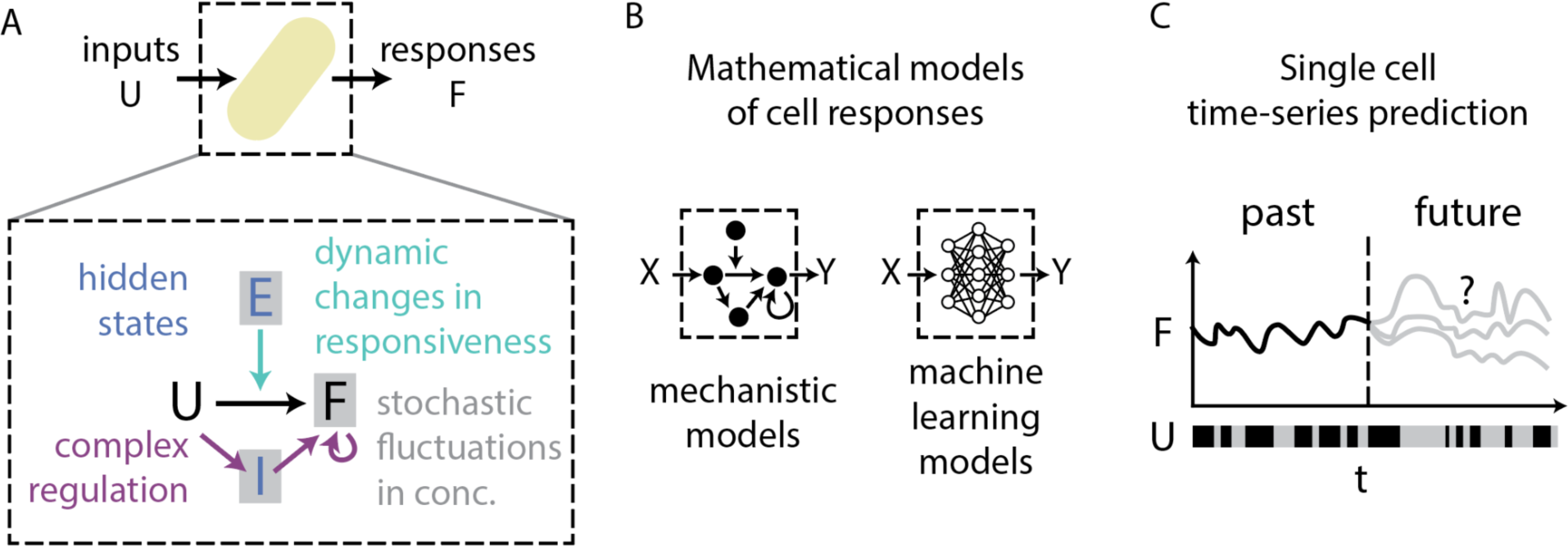
Machine learning models are well-suited to performing single cell time-series prediction. (A) The relationship between a time-varying input (*U*) and a cell’s dynamic response (*F*) is mediated by many phenomena. Stochasticity in gene expression can create fluctuations in concentration; globally shared resources also vary, creating extrinsic noise (*E*). The relationship between *U* and *F* may also be subject to complex regulation, and aspects of the cell’s state may be hidden, for example due to intermediate steps (*I*). (B) Mathematical models can be used to relate dynamic inputs and outputs. Mechanistic models explicitly account for many of the phenomena relating these inputs and outputs, while machine learning models can approximate these relationships with neural networks and do not require knowledge of the mechanism. (C) The ability of machine learning models to learn generic input-output relationships makes them well-suited to time-series prediction, where the future response of a cell to a particular input is predicted, using information about its past history of responsiveness.

Despite these challenges, various modeling approaches have produced informative and useful predictions of cell dynamics (Figure 1B). Ordinary differential equations (ODEs) represent the overwhelming method of choice for modeling how cells change with time. Even the earliest synthetic biology models^1,2^ showed the power of ODEs for explaining expression dynamics in terms of gene network architecture, and this continues to be the case^4^. Moreover, ODE models can be modified to explicitly incorporate the stochasticity of reactions^19,20^ or sources of single cell variability^21^, and can even be expanded to include the majority of reactions in a cell^22,23^. In the absence of complete information about a process, qualitative approaches, such as logic or Boolean models, can also be constructed. Such models can capture dynamics of gene regulatory networks, differentiation, and signaling pathways^24–27^. Markov models, which model transitions between specific cell states, can also provide useful descriptions of time-series data, such as single cell epigenetic dynamics^28^.

While these classical approaches to modeling and predicting cell dynamics are powerful, they require existing knowledge of the system, to either select an appropriate model structure or to identify reasonable reaction rates. However, in many cases, the parameters describing the system behavior are incompletely known or are poorly constrained by existing data^29–33^, or worse, the system itself may not be fully identified^34–36^. These challenges can be addressed with ensemble models^30,37^ or optimal design of experiments^12,38^, but these approaches require significant effort for each new model.

Machine learning provides an alternative to these bespoke methods. Machine learning models can flexibly capture a variety of input-output relationships, as demonstrated by their power to perform diverse prediction tasks across a wide array of fields^39–42^, including biology^38,43–46^. In recent years, machine learning approaches have been applied to predict biological time-series data, including p53 oscillations^47^, metabolic pathway dynamics^48^, responses to chemotherapy^49^, cell signaling^50^, and gut microbiome dynamics^51^. While these “black-box” approaches can be difficult to interpret physically or to extend to perturbations not present in the training data, such models are still incredibly useful. For example, machine learning can provide a faster alternative to mechanistic models to accelerate discovery^52^. More importantly, training such models provides an opportunity to find relationships between features where current mechanistic knowledge is absent.

In this study, we used a machine learning model to predict gene expression time-series data in single cells and performed a series of “stress tests’’ to understand the limits of prediction accuracy, ultimately finding our trained models to be highly capable of predicting future system responses given past data under experimentally realistic conditions. The general goal of our approach is to forecast how a cell will respond to a future series of dynamic inputs based on information about its past response dynamics (Figure 1C). In this way, the neural network predicts not just population average responses to dynamic inputs, but, given a single cell’s history, its specific response to any future input. As a representative example, we focused on a deep neural network architecture that we implemented in previous experimental work to predict optogenetically induced gene expression in single *Escherichia coli* cells^53^. Optogenetic systems are ideal test cases for predicting cell responses with machine learning because generating many examples of single cell responses to diverse inputs is experimentally feasible. Further, using microfluidic devices it is possible to image single cells for hours to days, generating substantial temporal information about each single cell^53,54^. Nonetheless, the neural network architecture developed here is expected to generalize to many other types of cell responses.

To understand how the quality of these predictions varies with noise, we simulated gene expression traces with measurement noise and various sources of stochasticity, and evaluated how well the trained predictor performed. Measurement noise produced a constant prediction error, whereas the stochasticity of the biochemical reactions added a small error that increased with time, reflecting increasing uncertainty about the cell’s distant future. Surprisingly, adding a variable for extrinsic cell responsiveness, whose dynamics affected the cell response but were not light dependent, did not increase the median error. This is an encouraging finding, as cells exhibit fluctuating levels of responsiveness in realistic experimental conditions. We also found that while accurately predicting the future required a minimum amount of information about the cell’s past, training models to predict even further into the future did not change their near term prediction quality. We also tested alternative genetic circuit structures. The addition of a hidden intermediate in a genetic circuit cascade did not significantly increase the prediction error relative to a simple activation case, but the delays introduced by the cascade increased the requirement for past information. Finally, we tested the ability of this neural network architecture to predict multimodal responses by training on examples of an auto-activation circuit. However, this approach minimized prediction error by predicting the average of the two possible bimodal outcomes, as in many cases both futures were equally likely. To address this, we implemented an alternative neural network architecture to predict the full distribution of possible future cell states, and found that it could correctly predict the bimodal outcomes of the auto-activation circuit. Together, these results show the power of deep neural networks for inferring complex, single cell responses in the midst of various types of noise across genetic circuit structures. Overall, the flexibility of this approach allows it to be extended to more challenging prediction tasks, such as predicting the full set of a cell’s possible futures. Such methods can enable many important goals in synthetic biology that rely on predictions, such as exploration of genetic circuit designs and precise control of gene expression.

## Results

### Machine learning models can infer response dynamics with different types of noise

Various sources of noise make it challenging to predict future cell responses (Figure 1A). Some of these, such as partially observed states or interactions between components, produce complex input-output relationships that can be inferred by analyzing past responses. Others, such as measurement noise or the stochasticity of biochemical reactions themselves, may make identifying the patterns of the input-output relationship more difficult, in addition to adding fundamentally unpredictable randomness. We therefore sought to quantify the impact of different sources of noise on how well a deep neural network can learn input-output relationships.

We focused on the behavior of the CcaSR optogenetic system activating a fluorescent reporter gene^3,55^ (Figure 2A). This optogenetic circuit is attractive because it features few steps between the input stimulus and the cell’s response, providing a simple initial test case. In this circuit, the addition of green or red light respectively promotes or stops the assembly of transcriptional machinery, thus increasing or decreasing the amount of reporter. Also, empirical responses of this circuit were already fit to an ODE model^56^, providing equations and parameters for generating synthetic data (Supplementary Methods). We used this as a basis to develop deterministic and stochastic models of this circuit where green light, after a brief 12 minute delay to approximate dynamics of phosphorylation and dimerization, linearly activates a constitutively expressed dimer, which then activates a downstream reporter with a Hill function response. The disassembly of the dimer (*H* in Figure 2B) by red light has the same delay, to capture dynamics of dephosphorylation and unbinding.

**Figure 2:**
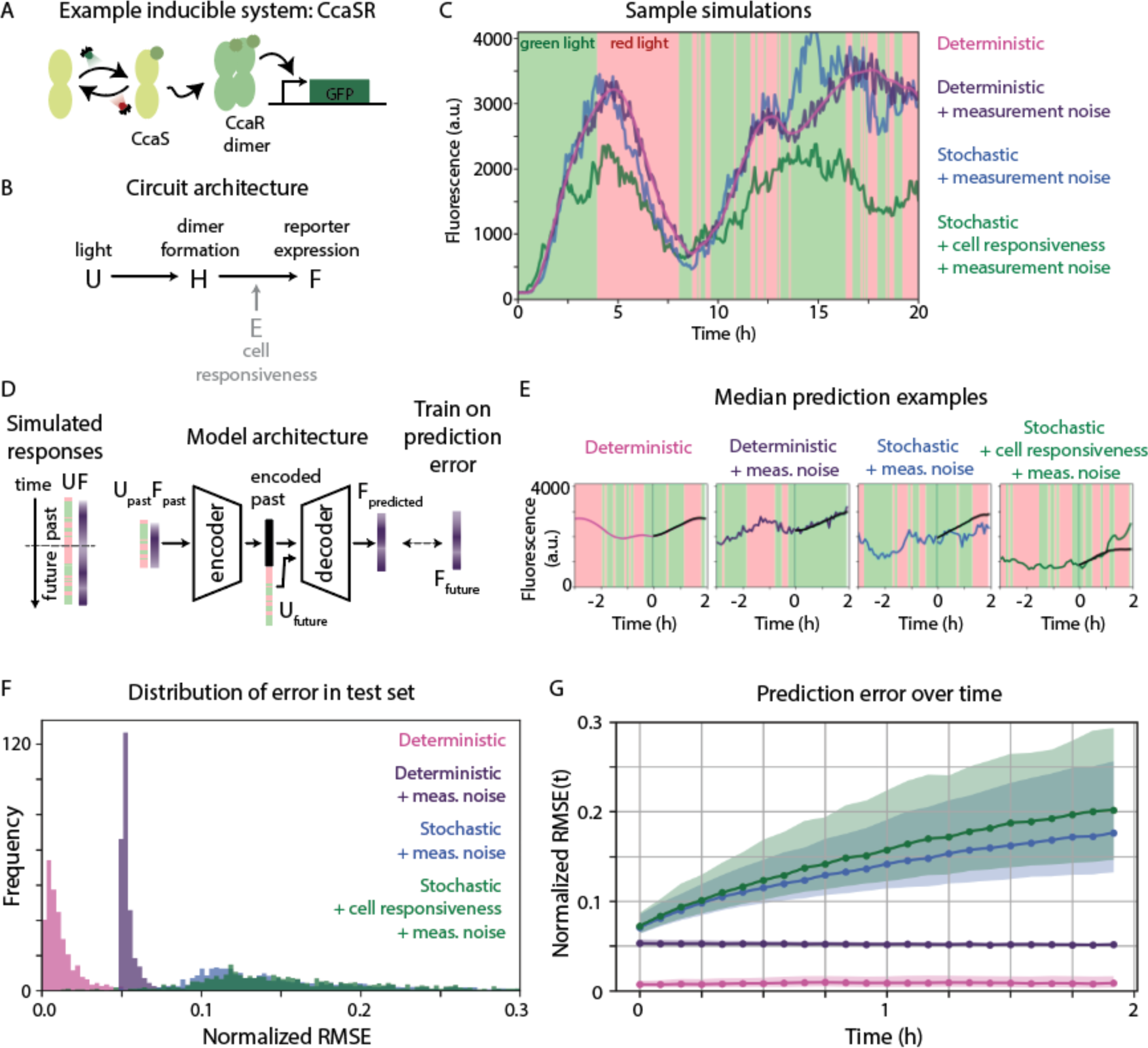
The deep neural network architecture can infer single cell response dynamics in the presence of multiple types of noise. (A) Simulations focus on a simple optogenetic activation circuit as a representative example. In the CcaSR system, green light activates CcaS, which activates CcaR to catalyze the formation of CcaR dimers that drive the expression of a reporter. Red light leads to reporter dilution by deactivating CcaS. (B) We simulated a simplified version of the CcaSR system, where light (*U*) activates a constitutively expressed dimer (*H*) that drives expression of a fluorescent reporter (*F*). In some simulations, we included another dynamic variable *E*, that captures light-independent changes in cell responsiveness. (C) Simulated responses to the same light input sequence with different types of noise show variability in dynamics between the four simulated datasets. (D) To train the model, simulated single cell responses are artificially partitioned into the past and the future. The model takes as inputs the past light sequence and associated cell response, as well as the future light sequence. An LSTM deep neural network encodes this as a latent representation of the cell’s past, which is concatenated with the future light sequence and passed to a decoder. An MLP decoder then predicts the cell’s response. The model is trained by reducing the error between the predicted future and the “ground truth” future, here taken from the simulations. (E) Predicted cell responses for each of the four noise models are shown with a corresponding example from the test set. Colored lines show the “ground truth” simulation, with one possible realization of the cell’s future. Black lines show the model’s predicted future for that cell and that light sequence. These are median examples of prediction quality from the test set. (F) The full distribution of error in the test set for each of the four noise conditions. As described in the main text, we compute the normalized root mean square error (normalized RMSE) for each example in the test set by computing the squared error between the predicted future and the 1000 possible realizations for a cell’s future, normalized to the simulated realization, and then averaging across the entire prediction horizon. (G) The normalized RMSE of each of the 1000 cells in the test set computed at each timepoint. Dots show median normalized RMSE, while the shaded region spans the 25th to 75th percentile. The colors correspond to the same four models shown throughout the figure.

Simulating cell responses allowed us to directly add or remove sources of noise and quantify their effects. We generated four sets of synthetic training data, adding new sources of noise to each dataset (Figure 2B). First, we considered data generated by deterministic ODEs. Even in the absence of noise, learning to predict these responses requires inferring a non-trivial response function, including the delay in activation and the accumulation of an unobserved dimer that non-linearly activates output. Second, we added multiplicative Gaussian measurement noise, which represents variation introduced as part of the measurement process, such as sCMOS camera noise or image segmentation errors. Measurement noise can complicate the inference and prediction of genetic circuit dynamics by obscuring the true state of the cell. Third, rather than deterministically evaluating the ODEs, we used the Gillespie stochastic simulation algorithm^19^ to evaluate the equations describing the cell response, and continued to add measurement noise. Lastly, we added a new variable and set of reactions to capture light-independent extrinsic fluctuations in cell responsiveness, reminiscent of cell aging^54^ or fluctuations in components for transcription and translation^15^ that can alter the dynamics of a circuit. These extrinsic fluctuations are simulated stochastically alongside the light activation dynamics, with the additional measurement noise. Besides increasing responsible variability, this new hidden variable (*E* in Figure 2B) adds a complexity to the cell response that can potentially also be inferred by the trained model, analogous to the inference of the process dynamics itself.

To demonstrate the effects of these sources of noise on cell responses, we simulated responses to the same light stimulus for each of the four cases (Figure 2C). While the deterministic responses are smooth in time, measurement noise creates variation around those responses. Stochasticity produces even more varied responses, which are further exacerbated by changes in the extrinsic cell responsiveness. For our synthetic training sets, we generated 10,000 examples of cells’ responses to different random light inputs for each of the four noise scenarios. This provides examples of not only how each system responds to light, but also how those responses vary across light durations and different historical contexts.

For each synthetic training set, we fit a deep neural network to predict cell responses (Figure 2D). At a high level of abstraction, this architecture takes as inputs the past light stimulus, past cell response, and a proposed future light stimulus, and predicts the most likely future response. To do this, the model first uses an encoder to identify key patterns in a cell’s past response, and then combines this with the proposed future stimulus to predict that particular cell’s future response. To begin, we used the same implementation as in our previous experimental work^53^, using a long short-term memory network (LSTM) for the encoder, which can flexibly take any length of past responses, and a multi-layer perceptron (MLP) as the decoder, due to its speed and ability to be parallelized for applications that optimize future responses (Methods). Note that this network architecture could be trained in a generative framework, where it would learn to generate a single possible stochastic realization of the future response. However, this would likely require hundreds of model runs to obtain a sufficient statistical understanding of a future response, thus we elected to use a straightforward supervised learning framework to directly output the most likely fluorescence level. We trained each predictive model with synthetic data, which provides many examples of past and future responses. We used 90% of the data for training and 10% for validation, keeping the model with the lowest error in the validation set. Note that throughout the manuscript, the word “model” refers to a neural network architecture trained on a particular dataset; thus, our analysis generated many “models,” all of which use this same architecture, unless otherwise noted.

To evaluate the quality of the prediction, we used our trained models to predict cell responses in a held-out test dataset. These test sets contained 1000 additional samples of light stimuli and simulated cell responses, none of which the trained models had seen before. We note that because we generated the randomized light inputs and simulated gene circuit dynamics in the same way for both the training and test sets, the range of fluorescence values in the training and test sets were largely overlapping (Methods, Figure S1A). As expected, the trained models’ predictions of the future approximated the “ground truth” simulated future with varying degrees of accuracy (Figure 2E). However, since many possible future responses are consistent with a cell’s past, a better measure of prediction quality would be a comparison of the predicted future to an ensemble of the possible cell responses (Figure S1B). We therefore generated 1000 possible future realizations for each cell in each test set. To quantify the prediction error relative to this ensemble, we calculated the normalized root mean square error (RMSE), the squared error between the predicted and simulated future, normalized by the size of the response, averaged across all possible futures. In this way, the values of RMSE can be interpreted similarly to percent error, with a value of 0.1 indicating a prediction error that is 10% of the actual value.

Specifically, for each of the 1000 cells in the held-out test set, we compute the following:

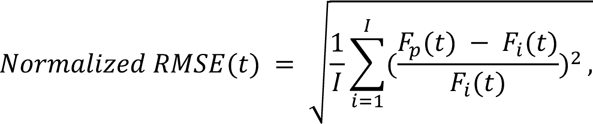

where *i* ∈ ⟦1, *I*⟧ indicates each simulated future (here *I* = 1000 for each cell in the test set). *F_p_(t)* is the predicted response at future timepoint *t*, whereas *F_i_(t)* is the *i*^th^ simulated future response for the cell. Because this error metric is defined at each point in time of the future prediction horizon, we compute average prediction quality by taking the mean across timepoints:

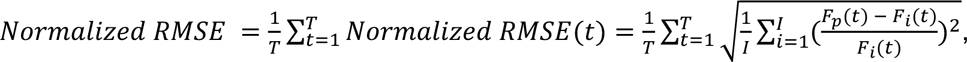

where *t* ∈ ⟦1, *T*⟧ indexes future timepoints.

The distribution of normalized RMSE across the test set for the different models showed that increasing noise corresponded to lower prediction fidelity (Figure 2F). Models trained on deterministic examples predicted simulated, deterministic responses with very low error, showing that the network architecture could correctly infer the response dynamics. By contrast, the addition of measurement noise introduced a minimum prediction error of approximately 5%, which is very close to the standard deviation of the sampled Gaussian noise. However, this error is not the result of incorrectly inferring response dynamics from the noisy examples; the model trained on examples with measurement noise predicted the average future response (i.e. the underlying deterministic dynamics) nearly as well as the model trained on deterministic data (Figure S1C). Rather, the smooth prediction cannot imitate the jagged jumps in the data produced by the fundamentally unpredictable measurement noise, creating a systematic and time-invariant error (Figure 2G).

The models trained on stochastic responses predicted the average response of each cell nearly as well as models trained on deterministic responses (Figure S1C). This suggests that the deep neural network can not only infer the underlying dynamics of the system from noisy examples, but is also able to perform state estimation for stochastically varying components. While the median and spread of normalized RMSE (cp. Figures 2F and S1C) were larger, this is consistent with the impossibility of predicting all aspects of the stochastic dynamics, where an increasingly large set of futures are equally possible (Figure S1B). Thus, even though the normalized RMSE increases with time as the spread of possible futures grows (Figure 2G), the model trained on stochastic responses (without changes in cell responsiveness) predicts the average dynamics consistently well across time (Figure S1D). Though the addition of changing cell responsiveness produced the hardest to predict dynamics, the prediction error for all models at early timepoints was not much larger than the case where only measurement noise was present (Figure 2G).

To benchmark this result against a method without a neural network, we also used an ODE model with a Kalman filter-based approach to predict the results of our stochastic simulations with time-varying cell responsiveness (Figure S2). Surprisingly, even though this ODE-based model has perfect knowledge of the system’s dynamics and parameters, our deep neural network is more accurate. Only if we directly provide information about the system’s state to the Kalman filter can it predict as well as the LSTM-MLP deep learning architecture. This is likely because Kalman filters are sub-optimal if system or measurement noise are not Gaussian, which is typically the case in single cell gene expression data and in our simulations in particular (Supplementary Methods).

Overall, this suggests that our LSTM-MLP architecture can infer deterministic dynamics very well, even if presented with noisy examples. For more variable stochastic dynamics, trained models can also predict average dynamics accurately, with only a small increase in percentage error. In general, predictions were best at short timescales, before the inherent randomness of the response accumulates into very large differences between equally likely futures.

### Prediction quality depends on training set size

For deep neural networks to correctly infer response dynamics, they require access to a sufficient number of examples. However, generating many long duration examples of single cell responses can be experimentally challenging for certain cell stimuli and responses of interest. Therefore, a critical question for the broad applicability of this approach is what amount of data is sufficient to allow the predictive model to generalize. To explore this, we focused on models trained on the stochastic circuit that also included changes in cell responsiveness and measurement noise, as we considered this the most biologically relevant (Figure 3A, green in Figure 2).

**Figure 3:**
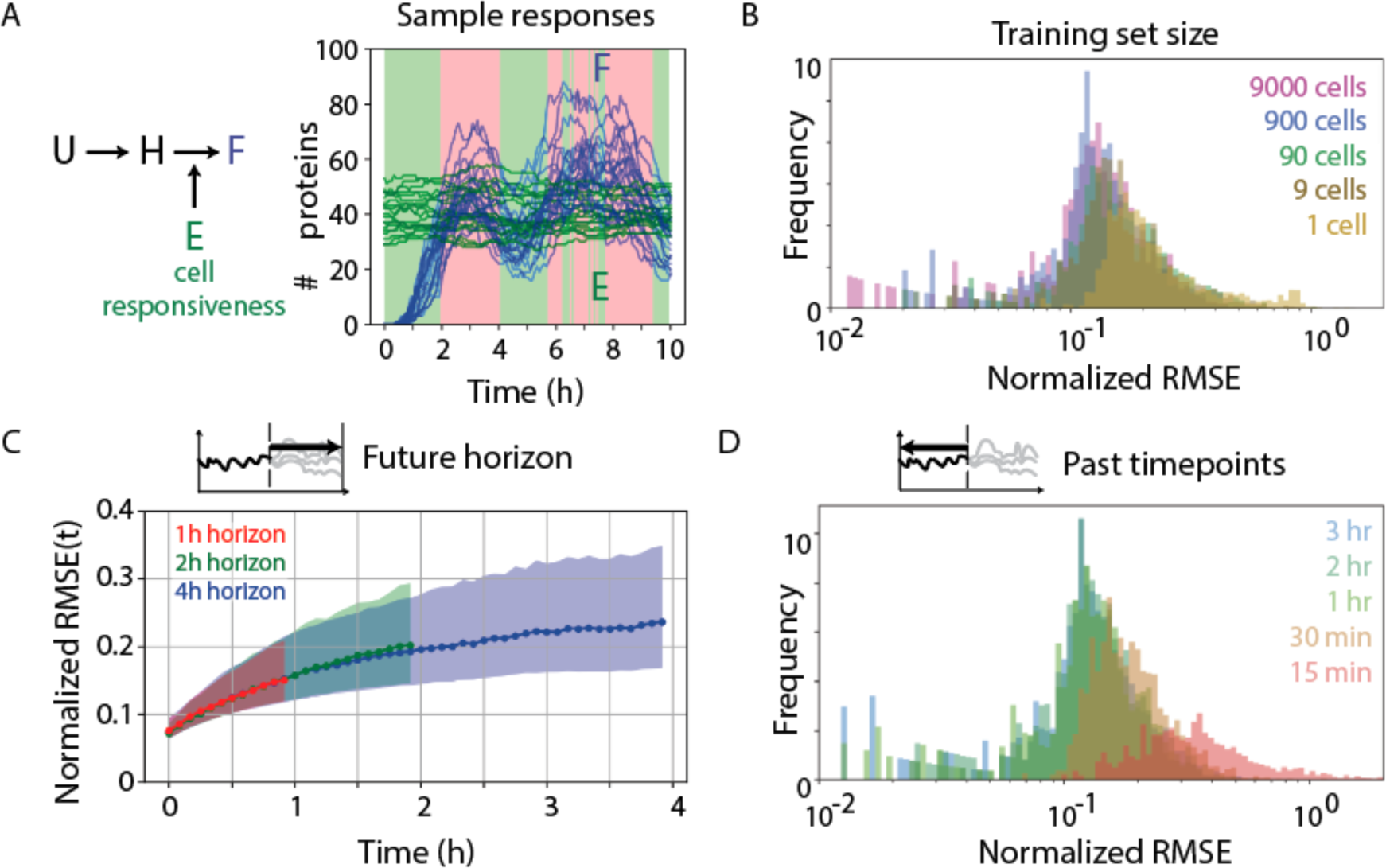
Larger training set sizes, more information about the past, and shorter prediction horizons increase prediction quality. (A) In the simple activation circuit, the overall level of cell responsiveness (*E*) tunes the maximal activation rate of *F.* Twenty sample responses of cells to the same light input are shown. At the initial timepoint, all species are absent, except for *E* which is randomly sampled. *E* (green) varies in time, but *F* (blue) varies both in response to light and changes in *E*. (B) The LSTM-MLP network architecture was trained using different training set sizes (neglecting validation data) to produce five different predictive models. All models used 3 hours of past information to predict a 2 hour horizon. The normalized RMSE is computed using the same 1000 example test set for all models. Median error increases as the training set size is reduced. (C) Three models were trained to predict responses that extended 1, 2, or 4 hours into the future, using 3 hours of the past and the full 10,000 example training set. Normalized RMSE(t) increases with time in the same way for all models, regardless of how far in the future they were trained to predict. (D) Five models were trained to predict the future using different amounts of the past data, a model hyperparameter that determines the input size. The original training set size of 10,000 cells and future horizon of 2 hours was used. The median and spread of normalized RMSE in the test set increased considerably when fewer than 30 minutes were used to predict the future.

We first trained models on different numbers of single cell responses, ranging from 9000 to 1. Each example response was 36h of randomized light stimuli and the simulated reporter response, sampled every 5 minutes. During every training epoch, the training data were sampled in batches, which included slicing out a particular 5h instance of the past and future from the longer 36h response. We first ignored the validation set and focused only on models that minimized error in the training set. When evaluated on the same 1000 example test set, models trained on either 9000 or 900 examples produced similar prediction errors (Figure 3B), suggesting that even a smaller dataset than we had initially used would have been sufficient. Only a 100-fold decrease to a training set with 90 examples increased prediction error.

We hypothesized that the poor generalization to the test set for smaller training set sizes was due to overfitting. In such cases, holding out some of the training data as a validation set can identify at which point in the training began to overfit the data. Indeed, in cases with only 9 or 90 training examples, the error in the held out validation set initially decreased as the training improved the weights, but then increased (Figure S3A). Keeping the model that performed best on the validation set improved predictions overall; considering 10 validation examples when training on 90 examples improved error in the test set to match models trained on 900 examples alone (Figure S3B). This suggests that holding out validation sets may improve prediction performance when training on very few examples.

The fact that models trained using 100 examples total can predict future responses as well as those trained on 9000 examples is somewhat surprising. However, this is likely due to the fact that a 36h recording of a single cell’s response actually contains many examples of 2h futures that follow 3h of the past. This suggests that for shorter recordings of cell responses, more total responses may be required to train the model. We therefore trained models where each “example” response was half as long (18h) or was precisely as long as the past and future horizon used for the prediction (5h). Without a validation set, these shorter duration responses required even more examples to match the best performance for 36h examples (Figure S3B). However, with a validation set and sufficient examples, even the shortest duration examples could be used to match the best performance when training on longer duration responses. All together, this suggests that longer duration example responses can partially compensate for a small number of single cell responses, though validation sets can also compensate for shorter recordings. However, this apparent tradeoff between number of cells and length of response may be particular to our synthetic data, in which the differences between early and late responses and between individual cells are largely quantitative differences in cell state. In real datasets, circuit performance may qualitatively differ between cells, either due to aging or spontaneous mutations. Moreover, for circuits that produce more diverse input-output relationships, the absolute training set size is likely different.

### Prediction quality depends on amount of past information, but not length of horizon

Because the LSTM-MLP deep learning architecture can flexibly accommodate different amounts of past and future data when making predictions, we were curious how tuning these values affected overall prediction quality. First, we considered the future horizon. The evolution of error with time (Figure 2G) suggests that longer term futures are harder to predict. We wondered if models trained to predict longer future horizons might sacrifice short-term prediction quality to predict longer term responses. We returned to the full 10,000 example training set, with 90% reserved for training and 10% for validation, and trained three models to predict either 1, 2, or 4h into the future given the cell’s most recent 3h of past responses. Across all prediction horizons, the normalized RMSE was the same at short timescales, and the median and spread of prediction error increased in the same way along the horizon (Figure 3C). This suggests there is not necessarily a trade-off in the prediction of short- and long-term futures, even though processes with different dynamics (e.g. fast activation of the dimer *H* relative to slower fluctuations in *E*) occur at the same time. Indeed, changes to the dynamics of *E* produced no change in prediction error when predicting over either a short (1h) or long (4h) horizon (Figure S4A,B).

We next considered the effect of changing how much of the cell’s past is provided as an input to the model. The LSTM-MLP architecture uses the entire cell’s past light stimulus (*U*) and responses (*F*) to build an encoded representation of the cell’s history, presumably summarizing features such as the unobserved concentration of dimer (*H*), as well as the degree and instantaneous rate of change in the cell’s responsiveness (*E*). In practice, long time series of past behavior are not always available, but using very short amounts of the past response runs the risk of undersampling these noisy dynamics. To assess the impact of how data about the cell’s past impacts prediction accuracy, we trained models to predict the same 2h future with access to different amounts of past information.

The number of past timepoints could be reduced from 36 (3 hours) to 12 (1 hour) without any change in normalized RMSE (Figure 3D). However, prediction quality decreased when only 6 timepoints (30 minutes) were used and deteriorated dramatically when only 3 timepoints (15 minutes) were used. We hypothesize that this cutoff reflects the number of observations or the absolute amount of elapsed time necessary to perform state estimation, either on the amount of dimer (*H*) or the light-independent changes in cell responsiveness (*E*). Considering that our model includes a 12 minute delay between when the light is changed and when it starts to have an effect, any architecture will likely struggle to accurately infer the cell’s responsiveness with only 15 minutes of past behavior. In fact, it may become easier to predict longer term futures, where there is more information about the historical light context. This would reverse what we have previously observed, where prediction error is initially low and grows over time. We indeed found that in the limit of a few past timepoints, the prediction of the initial response was no better than the prediction of the response endpoint, a trend which held across different dynamics for *E* (Figure S4C).

Together, these results suggest that a minimum amount of the past is required to make good predictions of the future and that, for the circuit studied here, this requirement is not set by the extrinsic noise, but rather depends on dynamics of the response to light, measurement noise, and stochasticity. Overall, with sufficient information about the past, models can be trained to predict arbitrarily far into the future, though the inherent unpredictability of the process produces a growing error that can eventually render those predictions moot.

### Cascade-associated delays increase need for past data

An advantage of machine learning models over mechanistic ones is their ability to predict input-output relationships without explicit knowledge of intermediate processes. Therefore, to explore whether our LSTM-MLP architecture could model more complex circuits, we simulated a cascade, a common motif in gene regulatory networks, by inserting an intermediate component downstream of the light-activated transcription factor and upstream of the reporter (Figure 4A). Because of this additional intermediate, the reporter input has a more complex dependence on the light as well as additional delays; the additional unobserved component may also make state estimation more complicated.

**Figure 4:**
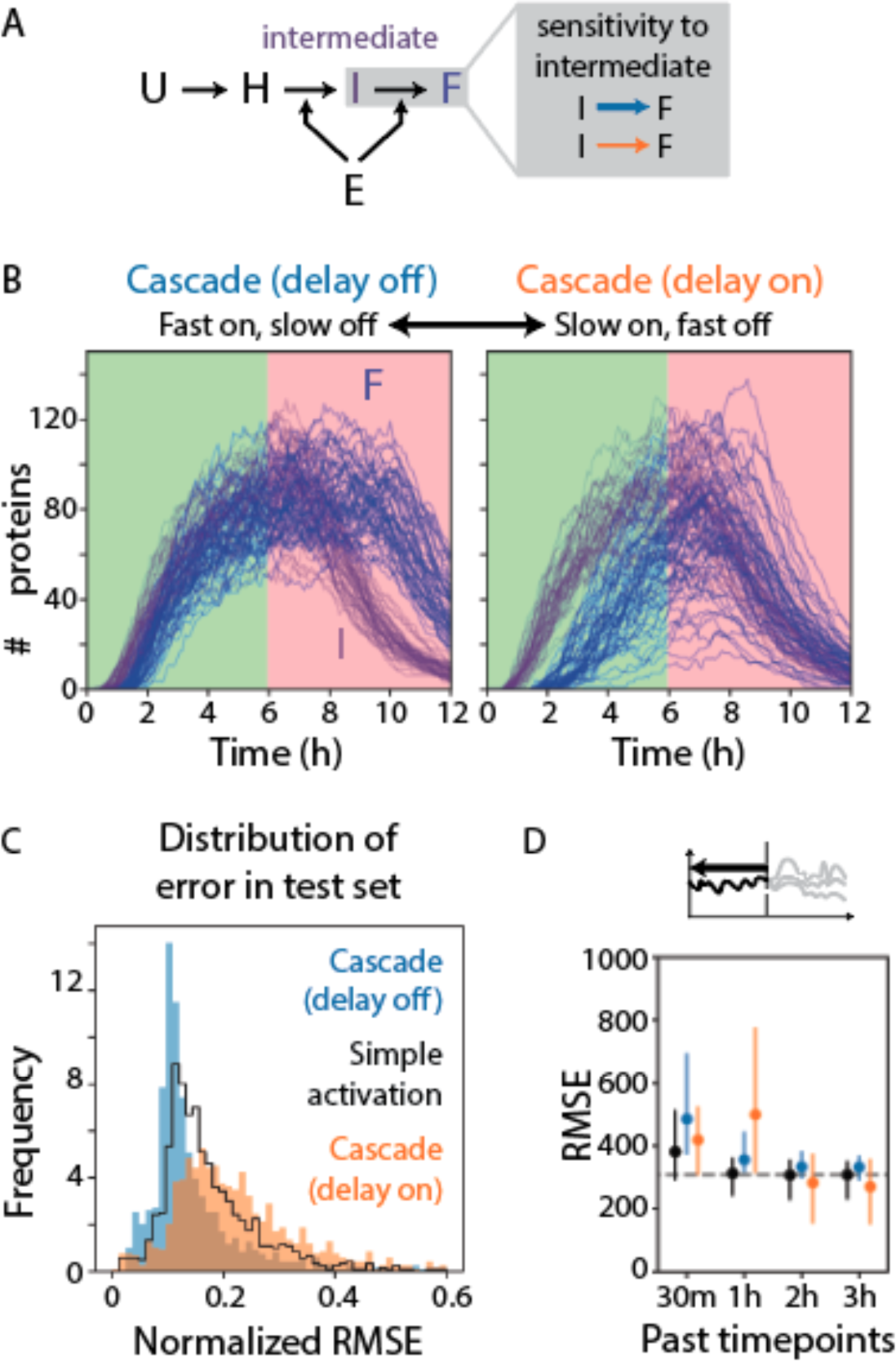
Delays in activation affect prediction quality more than delays in degradation. (A) In the cascade circuit, light (*U*) activates the constitutively expressed dimer (*H*), which activates an intermediate (*I*) that then activates a fluorescent reporter (*F*). The time-varying cell responsiveness affects both the expression of the intermediate and the reporter. By varying the amount of *I* required to half-maximally activate *F*, we simulated two versions of the cascade with different delays. (B) Fifty sample responses of *I* (purple) and *F* (blue) are plotted in response to a simple light input sequence to show their on and off dynamics. Though *F* approaches the same steady-state value for both circuits, the “delay on” circuit takes longer to reach it. (C) Two models were trained on 10,000 examples of either of the two cascade versions shown in panel B. Normalized RMSE was computed across the 1000 examples in the test set. Normalized RMSE for the simple activation circuit is reproduced from Figure 2F for comparison. (D) Additional models were trained to predict cascade dynamics using different amounts of the past, ranging from 30 minutes to 3 hours. Because the different distribution of fluorescence values for the two cascades produces systematic differences in normalized RMSE, we show here the RMSE. The dots correspond to the median error, while the lines span the 25th to 75th percentiles. The simple activation case (black) is reproduced from Figure 3D. The dotted gray line indicates the error for the simple activation case with 36 past steps, as a reference for the best prediction with the most information about the past.

We explored two regimes for this circuit by tuning the amount of intermediate required for half maximal activation of the reporter. When very little intermediate is needed to activate the reporter, the reporter accumulates quickly after the addition of green light, but has a slower response to red light, as the intermediate must be diluted before the reporter can decay (Figure 4B). In the second regime, much more intermediate is required to activate the reporter, producing a delay in activation under green light, but a faster response after red light is added. We tuned the maximum activation rate of the reporter such that both systems had the same steady-state behavior (Methods).

To assess prediction quality on this more complex circuit, we followed a similar procedure as before, training the LSTM-MLP on a simulated training set and evaluating error in a simulated test set. Models trained to predict responses for the cascade circuits had very similar normalized RMSE relative to models trained on the “simple activation” case (Figure 4C, Figure S5), suggesting that this more “complex” circuit could also be predicted by the LSTM-MLP architecture. We also asked if the delayed dynamics of both cascades impacted the quality of future predictions. We trained models to predict 1, 2, and 4 hours into the future. To avoid the confounding effect of different magnitude responses of the different circuits, we used the RMSE to quantify absolute prediction error. On short time scales (≤1h), the absolute prediction errors were similar (Figure S6A), though slightly larger for circuits with higher responses (e.g. “delay off” cascade). By contrast, at longer timescales (≥2h), this relationship inverted and the “delay on” cascade had the largest absolute errors as well as the widest variance. This faster accumulation of error for the “delay on” circuit was the most notable difference in prediction quality we observed between the three circuits. We hypothesize that this is because the “delay on” cascade may produce a more diverse set of response dynamics than the “delay off” circuit (Figure 4B), making its responses harder to predict.

We then considered the amount of past information required to predict the future, as the delay between a light stimulus and the resulting change in gene expression means the future response depends on stimuli that occurred further in the past. Indeed, both circuits required more past information than the simple activation circuit (Figure 4D), with the “delay on” circuit requiring even more past information than the “delay off” circuit. For models trained on the least amount of past information (i.e. 30 minutes, or 6 timepoints), absolute errors were similar but appeared at different points in the future horizon (Figure S6B). The model trained to predict the “delay off” cascade more accurately predicted the short-term future, while the model trained to predict the “delay on” cascade more accurately predicted the distant future (Figure S6B,C). Thus, the delay in activation dynamics increased the amount of past information required to predict the future, which suggests that this limit is not necessarily set by the LSTM-MLP architecture but by the dynamics of the process itself.

Together, these results show that circuits with additional hidden components are not necessarily harder for the LSTM-MLP architecture to predict. Rather, the underlying dynamics of the process, which can depend on parameter choice as well as circuit structure, are also important. Changing a single parameter in our cascade circuit altered where the delay occurred (i.e. in response to red or green light) as well as the overall speed and magnitude of the response. Delays in the response and how quickly the circuit approaches steady state likely affect prediction quality in distinct ways. For example, a cascade with more components can have a many hour delay between the input and the response, requiring past information from many hours prior to identify the stimulus that produced an upcoming response. Alternatively, even a circuit that responds immediately to light may approach steady-state slowly (relative to the timescale of light stimulus) or with diverse dynamics (e.g. determined by a hidden variable), which decreases long-term prediction quality or a model’s ability to generalize across new examples.

### A convolutional decoder can predict the entire distribution of future cell responses

We next asked how our prediction approach performed when applied to circuits that produce a non-trivial dynamic response, such as bistability. Specifically, we evaluated our approach on a positive feedback loop, where the downstream gene *F* is activated by light via the dimer and also by itself (Figure 5A). This system displays a hysteretic cycle where, at intermediate levels of light stimulation, two stable equilibrium points can exist (Figure S7). We began by using our initial network architecture that uses an MLP decoder (Figure 5B) to predict cell response. In cases where the cell behavior is unimodal, this network is able to predict a trajectory that matches the response of most future realizations in our test set well (Figure 5C). However, when the possible future responses are bimodal, the network predicts a trajectory that splits the difference between the two modes to minimize the potential prediction error (Figure 5D). This is a poor prediction, because an intermediate trajectory is unlikely to occur since the system will migrate towards only one of the two equilibrium points, but which of these two will depend on the stochastic response. This represents an important limitation of the single-trajectory prediction method we have used so far. Moreover, even in the unimodal case, the prediction does not convey any information about the variability of the response to a particular light sequence, in some sense also making this a poor prediction of the true cell response.

**Figure 5:**
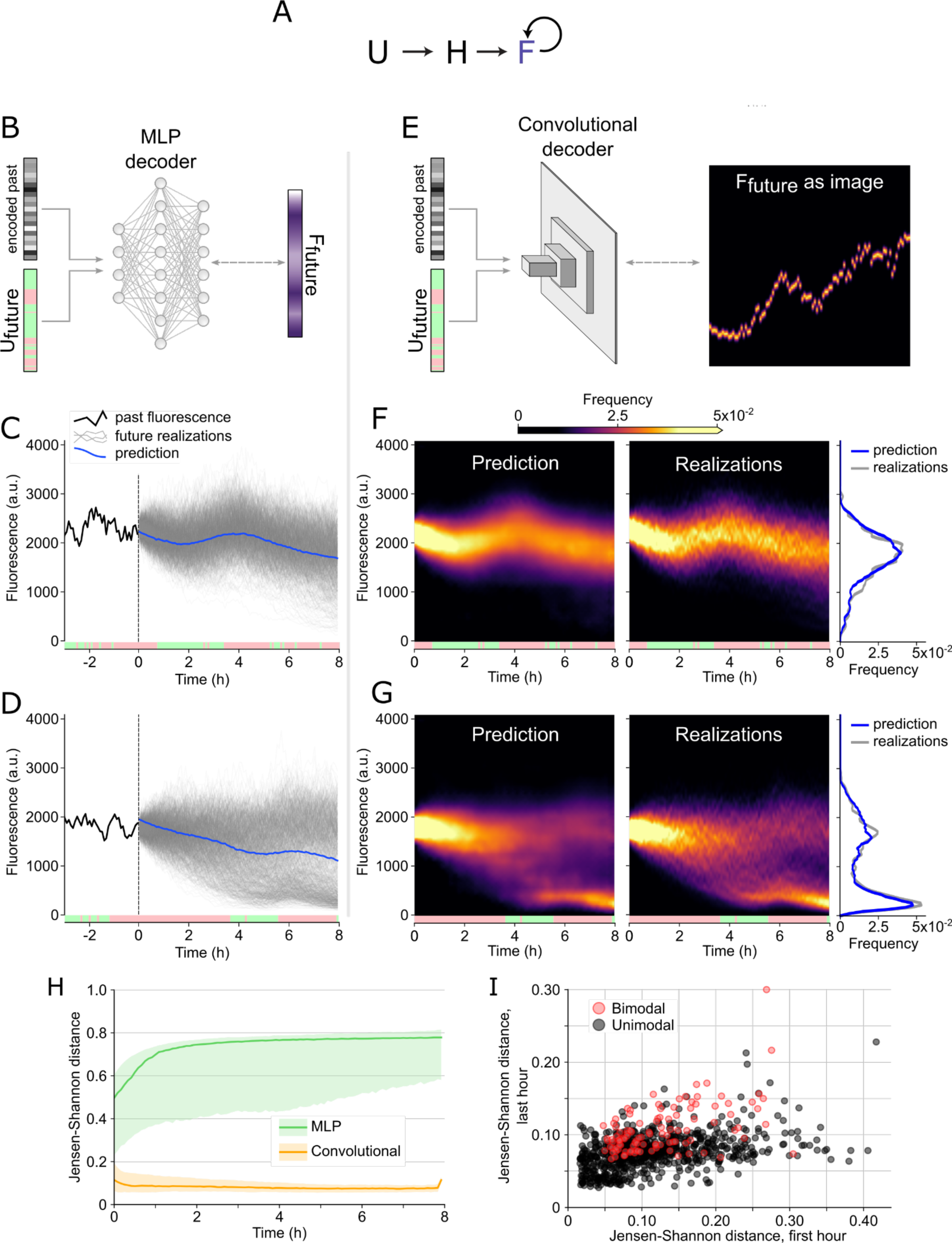
A convolutional decoder can predict cell response probability landscapes. (A) Schematic of the auto-activation circuit. Light *U* activates the dimerization of *H*, which in turn activates the reporter protein *F*. In addition, *F* also activates its own expression. (B) Schematic representation of the MLP decoder architecture. The final layer is trained against a vector representation of the future fluorescence response. (C) Test sample with unimodal response. The red and green light sequence is shown at the bottom of the plot. The past 3 hours of the cell’s response are shown as a solid black line. The vertical dashed line represents the separation between past timepoints, and the future response to predict. Thin gray lines represent the 1000 ground truth realizations of this simulated cell’s future response. The solid blue line represents the predicted single-trajectory response from the original neural network with a MLP decoder. (D) Test sample with bimodal response. In this sample, the prediction from the MLP decoder is unlikely but minimizes prediction error. (E) Schematic representation of the convolutional decoder architecture. The final layer is trained against an image representation of the future fluorescence response (Figure S8). (F) Prediction from the convolutional decoder for the unimodal sample. The prediction from the decoder approximates the actual probability landscape compiled from all 1000 future realizations from panel C. The distributions at the last timepoint for each are shown to the right. (G) Prediction from the convolutional decoder for the unimodal sample. Even in the bimodal case, the network minimizes loss by producing a prediction similar to the bimodal landscape from panel D. (H) Jensen-Shannon distance over an 8 hour horizon for the two network architectures. The green region represents the distance over the test set for the MLP decoder, while the orange region represents the convolutional decoder. Shaded regions represent the 25th-75th percentiles across the test set, and the solid lines show the median. The sharp increase in distance over the last few timepoints for the convolutional decoder may be due to edge effects in expanding convolutional networks. (I) Jensen-Shannon distance averaged over the first and last hour of the prediction horizon for unimodal (black) and bimodal (red) test sample cases.

To address this issue, we introduced a novel network architecture that uses an expanding convolutional decoder to predict cell responses as probability landscapes across time and fluorescence levels (Figure 5E). The convolutional decoder replaced the MLP decoder we implemented in all earlier designs (Figure 2D, 5B), but the architecture otherwise remained intact. When training this new network, we used exactly the same training data as with the previous network, using past time series of fluorescence and optogenetic events processed into a latent vector of fixed size, that is then concatenated with the future optogenetic sequence. The only difference is that the future single cell trajectory used as training ground truth was formatted into an “image” representation, with the horizontal and vertical coordinates corresponding to the time and fluorescence levels, respectively (Figure S8).

As with our previous prediction network, the decoder cannot learn to predict exactly how a single, stochastic cell will respond to future light events. However, by allowing it to represent the cell’s future as an “image,” with multiple fluorescence levels possible at a given point in time, the network minimizes the loss by predicting cell responses that closely match the full probability landscape of the potential cell responses. This is true not only for unimodal landscapes (Figure 5F), but also when the cell response is bimodal (Figure 5G). This is somewhat surprising given that the model is only trained on single realizations of a cell’s future, yet the model infers that these dynamics allow multiple possible futures. Thus, this new approach not only provides information about the variability in cell responses, it also provides an elegant approach to multimodal time-series forecasting.

To evaluate how similar the model predictions are to the actual probability landscapes in our test set, we computed the Jensen-Shannon distance^57^ between the distributions at every future predicted timepoint for both the convolutional decoder and the MLP decoder (Figure 5H). As expected, the prediction from the convolutional decoder is much closer to the actual probability landscape than the single-trajectory prediction from the MLP decoder. Interestingly, the distance for the convolutional decoder prediction decreases in the first 30 minutes and then remains constant across the prediction horizon. This may occur because at early timepoints the network faces a state estimation problem, where the response distribution depends strongly on the state of the cell right before the prediction. Between the stochasticity of the cell and measurement noise, a certain degree of uncertainty is inevitable, and this makes it challenging to predict how the cell distribution will look in the near future. However over the long term, the responses tend to depend less on the initial state of the cell and more on the illumination sequence, making the long term distribution of cell responses easier to predict.

One way to explore the impact of state estimation errors on prediction quality is to look at correlation between prediction error at early and late timepoints. We hypothesized that these should be correlated, and perhaps correlated more strongly in cases where stochasticity generates a bimodal response First, we classified the examples in the test set as either unimodal or bimodal with the statistical dip test^58^ (Figure S9). We split our dataset into 108 bimodal and 804 unimodal cases, with the remainder representing ambiguous cases. We found that the Jensen-Shannon distance was overall larger for bimodal samples (Figure 5I). We also found that a large Jensen-Shannon distance in the first hour of the prediction horizon was more strongly correlated with a large distance in the last hour for bimodal samples (Pearson correlation r=0.53, p<10^-8^) than for unimodal ones (r=0.39, p<10^-29^). These results indicate that not only are bimodal landscapes more difficult to predict accurately, but also that early errors in state estimation can lead to higher prediction error specifically in cases when the future cell response is more dependent on its past, which is particularly true for systems exhibiting hysteresis. Interestingly however, even for the worst prediction of our test set, the model accurately predicts the position of the distribution modes (Figure S10), and appears to only fail at predicting their relative frequency because of an erroneous initial state estimation.

These results show that by changing the decoder architecture, we can train models to predict the probability landscape of cell responses to dynamic inputs, even when the response is multimodal. Notably, even though the model is trained only on single realizations of a cell’s possible future, it learns how these dynamics enable diverse possible futures. This ability to track the full set of possible outcomes causes prediction error to decrease with time, the opposite of what we observed with the MLP decoder architecture, which could only predict one possible cell future.

## Discussion

Predicting cell responses is important for the design of new biological systems, but remains challenging in cases where the underlying process is stochastic or incompletely identified. Machine learning methods, which can model input-output relationships in the presence of noise and without mechanistic information, present an exciting alternative. Using computational datasets simulating the optogenetic control of gene expression in single cells, we explored the ability of a deep neural network to predict cell responses in the presence of different sources of noise and across different classes of genetic circuits. We found that our models could correctly infer the underlying dynamics of a cell response even in the presence of measurement noise and stochasticity in the biochemical reactions, though the latter resulted in accumulating error as the time series progressed. By contrast, adding additional components, such as variable cell responsiveness or an intermediate, did not increase error considerably. Rather, prediction quality depended strongly on the training set size and the amount of the cell’s past the model had access to when predicting future responses. Our stochastic simulations highlighted a fundamental challenge to predicting future responses. Requiring the model to predict a single future by definition increases the prediction error because the breadth of possible futures also tends to increase with time. We therefore implemented a novel approach to predict the full distribution of possible cell responses, which accurately predicted the future states of even a multistable circuit. Together, our work highlights the impressive potential of deep neural networks to infer cell dynamics in the presence of noise, and the flexibility of machine learning methods to perform diverse prediction tasks.

Extending this approach to a broader set of cell responses depends on many other important and interesting questions. For example, though we did vary the training set size, various other aspects of the training data, such as the overall structure or diversity of the light inputs, likely also affect the model’s ability to infer underlying dynamics. We also suspect that important quantitative relationships, such as the sampling rate relative to the speed of cell response, or the magnitude of the cell response relative to measurement noise, will also affect prediction quality. Models can also be trained on examples of more complex genetic circuits or even simulations of arbitrary transfer functions, in order to systematically probe the prediction capabilities of this architecture. These could be extended to processes with non-binary inputs, reflecting graded light intensity or concentrations of inducers. Moreover, possible variants of the encoder-decoder architecture have not been fully explored. An LSTM decoder could predict arbitrarily far into the future, whereas an updated encoder may perform better state estimation on smaller samples of the cell’s past. Generative approaches could also be used to rapidly provide possible future response realizations, to mirror the stochasticity and temporal statistics of single cell responses. Lastly, training models on simulated data of related cell processes could be used to explore the potential of transfer learning, where models trained on one dataset can be used to predict processes in another context, with little or no additional training.

Pursuing these questions can lay the groundwork for exciting future applications, most importantly predicting the responses of living cells across diverse inducible systems in natural and synthetic circuits. Deep learning methods are likely powerful and flexible enough to predict responses for many observable outputs to inducible inputs, giving them broad applications across processes such as metabolism and cell differentiation even in the absence of precise mechanistic information. Moreover, because any feature can be used as an output, these models could be used to infer relationships between variables that vary across scales, such as the effect of a signaling protein on cell morphology or size, which is not possible with most ODE models.^59^ When coupled with optimizers that identify the best possible input sequences to produce a desired output, such models can enable model-predictive-control of an equally broad set of biological variables.^53^ The unique abilities of machine learning models can make other prediction tasks more feasible than they would be with traditional ODE approaches. For example, predicting the distribution of future states may enable the inference of noise propagation and multimodality from single cell traces, even without an expert-built model. At the same time, deep learning methods can complement existing ODE approaches. Machine learning models can approximate ODE models, enabling fast and broad parameter exploration before validating results with the slower mechanistic models.^52^ Architectures such as the LSTM-CNN can also be used to approximate solutions to the chemical master equation, useful for not only estimating average behavior, but higher order moments.^60,61^ Future work may also overcome the “black box” nature of these models by reformulating them to have greater interpretability,^62,63^ revealing what features are necessary to predict future responses, or using other methods to infer symbolic relationships, guiding the formulation of a more mechanistic model.^64^ Overall, the power and flexibility of machine learning methods make them well-suited to a diverse set of prediction tasks that will enable future synthetic biology designs.

## Methods

### Code for simulations and deep learning models

All code necessary to simulate cell responses, generate training and test sets, implement and train the neural networks, and generate the figures in this manuscript is available in our deepcellcontrol repository on Gitlab, under the “simulations” branch: https://gitlab.com/dunloplab/deepcellcontrol/-/tree/simulations

### Cell simulations

To simulate the behavior of cells under time-varying optogenetic inputs, we wrote a custom implementation of the original version of the Gillespie algorithm^19^ in Python. This custom code made it easier for us to implement optogenetic events, delays, and to investigate system behavior. However, we also wrote a parallel implementation based on the GillesPy2 Python library (https://github.com/StochSS/GillesPy2) to verify the validity of our simulations. For the deterministic simulation, we wrote an ODE solver based on the SciPy library (https://scipy.org/) that used the same set of parameters and propensity functions. Values for all species were sampled every 5 simulated minutes, and the optogenetic inputs were applied at the same intervals, to mimic the conditions of our experimental platform^53^.

The solvers were implemented as methods to Python classes that simulated the behavior of the circuits shown in this study. Specifically, we created a main parent class featuring system propensity functions, parameters, and species that described the behavior of the simple circuit described below. This model was based on the model from Chait et al.^56^, with minor modifications to propensities to increase the stochasticity of the system, and to parameters to approximate the dynamics observed in experimental data from our previous work^53^. Then, we created child classes based on this model where the propensity functions, parameters, and species were altered to emulate the behavior of different circuits (see Supplementary Methods).

To simulate measurement error and to match the dynamic range of our experimental data, we multiplied simulated reporter protein numbers by a factor of 40 and added an offset of 100. We then clipped these “measured fluorescence” values to our microscope camera’s range of [0, 4095]. When measurement noise was applied, we added 5% Gaussian noise to these values before clipping. For measurement noise, we largely considered the errors associated with microscopy and downstream analysis. The noise associated with an sCMOS camera is largely Poisson distributed, related to the arrival of photons and generation of electrons, which becomes normal in the limit of a large characteristic rate^65^. Individual cell fluorescence values are typically extracted from microscopy images by segmentation, the possible errors of which we also approximate with a Gaussian. We ultimately chose to model noise with a multiplicative factor *M*∼Normal(μ = 1, σ = 0.05) because it produced traces similar to what we had observed in other experiments. The resulting fluorescence “measurements” are used in the training, validation, and test sets.

### Genetic circuits

We simulated the behavior of three main circuits. First, we simulated a “simple circuit” (Figure 2A,B) where light activates a constitutively expressed dimer that drives the expression of a reporter protein. In this simple case, green light activates expression of the reporter and red light deactivates the CcaSR system, which leads to reporter decay because of dilution. This circuit was simulated with the deterministic solver, with or without added “measurement noise.” It was also simulated with the stochastic solver, with added measurement noise. Finally, we added a “cell responsiveness” proxy molecule to the simulations under a Poisson process that evolves randomly over time and is independent of the light sequence, to approximate extrinsic cell-to-cell variability.

The second circuit is a cascade where the CcaSR system is used to activate the expression of an intermediate gene that in turn activates the expression of the reporter protein. The activation of the reporter by the intermediate follows a Hill function. To explore different versions of the cascade, we changed the parameter governing the number of intermediate molecules required for half maximal induction (i.e. *K_I_*, see Supplementary Methods). To ensure that all versions of the cascade had the same steady-state behavior and thus approximately similar fluorescence values, we adjusted the maximal activation rate for the reporter (Supplementary Methods).

Finally, the third circuit is an auto-activating loop where the CcaSR system is simulated to activate the reporter protein, but the reporter protein is also self-activated. We selected the parameters of this positive feedback loop such that the circuit displayed hysteretic behavior for intermediate values of average optogenetic inputs (Figure S7).

Detailed description of these simulation models including propensity functions and parameter values are provided in Supplementary Methods.

### Dataset generation

For each circuit, we generated training sets of simulated cell responses to random optogenetic sequences. Unless otherwise specified, 10,000 cells were simulated per training set, for 36 hour-long sequences to match what we were able to achieve experimentally^53^. The optogenetic sequences were randomly generated as described in our previous work^53^, by binarizing a one-dimensional random walk. These simulated training sets were saved to disk and used for network training. Training was done in batches, where in each batch, cells were uniformly sampled, and a uniformly random timepoint in each sequence was picked. The “past” time-series data prior to this point was fed as input to the encoder. The “future” optogenetic sequence over the prediction horizon was concatenated with the encoded representation of the past and fed into the decoder. Finally, the “future” fluorescence values were used as the ground truth for training. To evaluate the impact of smaller datasets, we sub-sampled the original training sets and saved these versions to disk for training.

We also generated test sets for each cell class to quantify the accuracy of our prediction networks after training. However these test sets differed from our training sets in one crucial way. Because our “cells” are simulated, we can compute several stochastic realizations of a cell’s response to an optogenetic sequence that are all consistent with its past. To do this, we first generated a single-realization of the response over a 24 hour-long optogenetic sequence for 1000 different cells. This constitutes the “past” part of our evaluation sets and serves as input to the model being evaluated. Then, for each of these 1000 samples, we computed 1000 realizations of the response to a “future” optogenetic sequence of 4 hours (or 8 hours for the auto-activation case, which allowed us to evaluate the probability landscape prediction accuracy over long periods and better show bistable responses). In this way, we estimated the distribution of possible responses to the future sequence, which we then used to compute prediction accuracy by comparing the prediction (either a single value, for the MLP decoder, or distribution of predictions, for the convolutional decoder) to the distribution of possible futures.

To train the convolutional decoder version of our network we used the same datasets as we did for the MLP decoder version, but re-formatted the “future” fluorescence ground truth trajectory as an image for training. A naive way to do so would be to represent fluorescence by the vertical pixel position in the image and time by the horizontal pixel position, setting the pixels corresponding to the time-series data to 1 and all remaining pixel values to 0. However, the dimensions of the “image” must not be too large or it slows down training. For technical reasons related to the network architecture, image dimensions also need to be multiples of 8. We therefore binned the [0; 4095] floating-point fluorescence values into 96 bins. To avoid losing too much information by directly quantizing fluorescence values, we applied a Gaussian kernel to these values, and the resulting Gaussian curve itself was binned into the corresponding pixels along the fluorescence axis (Figure S11). Thus, the actual fluorescence value can be reconstructed from the binned image representation with minimal loss of information. Along the time axis of the image, we simply sampled all timepoints, and because we use an 8 hour horizon with a 5 minute sampling rate, the time axis is also 96-elements long. Thus, the resulting image is a 96x96 representation of the single cell, single realization response to the “future” optogenetic sequence, that can then be used as the ground truth to compare to the output of the convolutional decoder in training. For the test set, we used the same approach to compute the “actual” probability landscape: For each sample, we computed the image representations of each of the 1000 future realizations of the cell response to the future optogenetic sequence, and then averaged all 1000 images together to obtain the probability landscape.

### Machine learning network architectures

All models were written with the Tensorflow library and the Keras API. The first network is almost identical to the one introduced in our previous work^53^, which features an encoder-decoder architecture (Figure S12). Briefly, the encoder contains two LSTM layers of 64 and 16 units that process past fluorescence and illumination time series into a 32-element latent vector representation. The decoder is a MLP that is trained to predict future fluorescence levels based on a concatenation of the latent representation of the past with the future optogenetic events. The decoder features 5 densely-connected layers of 32 neurons, with rectified linear activation functions. The final layer is a densely-connected layer of 12, 24, or 48 units depending on the prediction horizon (1, 2, or 4 hours, respectively), with a linear activation function. We used a multi-layer perceptron decoder because it can be parallelized more efficiently than another LSTM.

To predict probability distributions, we introduced a second architecture that replaces the MLP decoder with an expanding convolutional decoder, an architecture similar to the decoder part of variational autoencoders for image generation (Figure S13). The LSTM encoder architecture is unchanged from the first model, where the latent representation of the past time series is also concatenated with the future optogenetic time series. However, before being passed to the decoder, this vector is then used as input to a single densely connected layer and then reshaped into a 3D tensor of shape 12x12x16, and a 2D convolutional layer with 16 filters is applied. This 3D tensor is then fed through 3 up-scaling blocks composed of 2 layers: a transposed convolutional layer with a stride of 2 that doubles the size of the tensor along its first 2 dimensions, and then a convolutional layer to further process the result. The layers in these three consecutive blocks feature 32, 16, and 8 filters, and upscale the image by a total factor of 8, generating a final tensor of shape 96x96x8. This tensor is processed by a final convolutional layer into an output of shape 96x96x1, corresponding to our horizon of 96 timepoints and 96 quantized fluorescence levels.

### Training and evaluation

The networks were trained for 200 epochs, with a batch size of 100, and 200 steps per epoch. Unless otherwise specified, the training set was split into 90% training data and 10% validation data. Root mean square error (RMSE) and mean absolute error (MAE) were evaluated over the validation data after each training epoch to assess overfitting. After each epoch, model weights were saved to two different files depending on whether they minimized training loss or validation data error. Unless otherwise specified, the weights that had minimized the validation error were used after training. We used the Adam optimization algorithm^66^, with a learning rate of 0.001. For the networks with a MLP decoder, mean square error was used as training loss. For the network with a convolutional decoder, Huber loss was used^67^.

We evaluated the quality of our single-trajectory predictions by computing various error metrics. In most cases, we computed the normalized RMSE or normalized RMSE(t) across all the ground truth realizations for each sample in the test set, as defined in the main text. Briefly, for each sample, the error between each of the 1000 realizations and the prediction was computed, squared, and then the mean error across all realizations was computed at each timepoint, resulting in a single, time-distributed vector per sample in the test set. Finally, the square root of all elements in these vectors was computed. This way, we were able to not only obtain the normalized RMSE over time, but also a distribution of that error across the test set, which would not have been possible to compute without multiple realizations of future ground truths. In cases where comparing normalized RMSE was values was challenging, because the two test sets had systematically different ranges of fluorescence values and thus very different normalization values, we computed RMSE (or RMSE(t)) as follows:

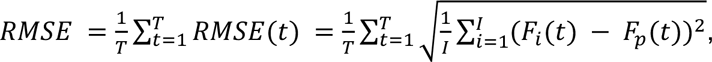

where *i* indexes the possible future realizations of a cell’s response, and *F_p_(t)* is the predicted response.

In many cases, the RMSE or normalized RMSE were large because the spread of possible values for *F_i_(t)* was also large and could not be approximated by a single future prediction. We were therefore curious if the model was correctly predicting the average future response, i.e. producing the minimal possible RMSE. We therefore also computed the normalized root mean squared error relative to the average future (NRMSE_average_) as follows:

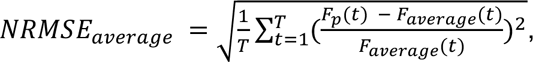

where *F_average_(t)* is the average future response, defined as:

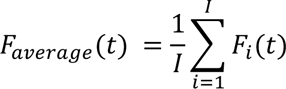

with *i* as the index for each cell’s possible future responses. To explore how this error evolves with time, we also computed a normalized absolute error:

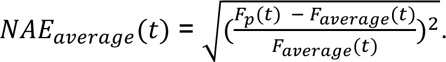

To evaluate the quality of our probability landscape predictions, we computed the Jensen-Shannon distance^57^ across time between the model predictions and the Gaussian kernel histograms of the test set ground truth described above. The Jensen-Shannon distance is a measure of the similarity between two probability distributions. It is calculated as the square root of the Jensen-Shannon divergence, which is a symmetric version of the Kullback-Leibler divergence, using the following formula:

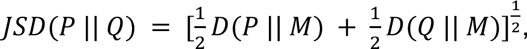

Where

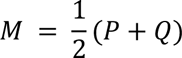

and *D*(*P* || *Q*) is the Kullback-Leibler divergence, computed as:

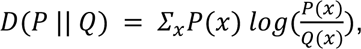

with *x* as the bins of fluorescence values used for the convolutional neural net prediction. By definition, the Kullback-Leibler divergence is not symmetric, as it quantifies the information lost by using one distribution to approximate another, which is not the same as its inverse. The Jensen-Shannon divergence involves computing the Kullback-Leibler divergences twice, between either of the two distributions and the average of these two distributions. The resulting Jensen-Shannon distance ranges from 0 to 1, where 0 means the two distributions are identical and 1 means they are completely dissimilar.

We used the statistical dip test^58^ to evaluate whether the distribution over the last hour of the evaluation histograms was unimodal or bimodal. This test works by calculating the empirical distribution function over all sample points, and then minimizes the maximum vertical difference between a unimodal reference distribution and the cumulative empirical distribution. This minimized difference is thus indicative of how close the empirical distribution is to being unimodal, and after analyzing our evaluation dataset with this test, we chose a dip difference of 3!10^-3^ as a threshold below which our data was considered unimodal, and 6!10^-3^ as a threshold above which our data was considered bimodal (Figure S9).

## Author Contributions

H.E.K. and J.B.L. conceived and designed the study with input from M.J.D. H.E.K. performed simulations and data analysis to compare noise models, analyze dataset requirements, and assess the impact of delays. J.B.L. developed the deep learning architecture and prediction method; he also developed and implemented the probability landscape approach. A.S.K. provided input on the results and analysis. H.E.K., J.B.L., and M.J.D. wrote the manuscript with input from A.S.K.

## Supporting Information

Supplementary figures and methods, including full model descriptions and their parameters, neural network model architectures, and additional analysis.

## Acknowledgements

We thank members of the Dunlop and Khalil labs for input. This study was supported by the National Science Foundation (NSF) grants MCB-2032357 and MCB-2143289 and the National Institutes of Health (NIH) grants R01AI102922 and R01EB029483. H.E.K. is a Damon Runyon Fellow supported by the Damon Runyon Cancer Research Foundation (DRG-2472-22).

## References

(1) Elowitz, M. B.; Leibler, S. A Synthetic Oscillatory Network of Transcriptional Regulators. Nature. 2000, pp 335–338. https://doi.org/10.1038/35002125.

(2) Gardner, T. S.; Cantor, C. R.; Collins, J. J. Construction of a Genetic Toggle Switch in Escherichia Coli. Nature. 2000, pp 339–342. https://doi.org/10.1038/35002131.

(3) Olson, E. J.; Hartsough, L. A.; Landry, B. P.; Shroff, R.; Tabor, J. J. Characterizing Bacterial Gene Circuit Dynamics with Optically Programmed Gene Expression Signals. Nat. Methods 2014, 11 (4), 449–455.

(4) Wilson, M. Z.; Ravindran, P. T.; Lim, W. A.; Toettcher, J. E. Tracing Information Flow from Erk to Target Gene Induction Reveals Mechanisms of Dynamic and Combinatorial Control. Mol. Cell 2017, 67 (5), 757–769.e5.

(5) Benzinger, D.; Ovinnikov, S.; Khammash, M. Synthetic Gene Networks Recapitulate Dynamic Signal Decoding and Differential Gene Expression. Cell Syst 2022, 13 (5), 353–364.e6.

(6) Villaverde, A. F.; Bongard, S.; Mauch, K.; Balsa-Canto, E.; Banga, J. R. Metabolic Engineering with Multi-Objective Optimization of Kinetic Models. J. Biotechnol. 2016, 222, 1–8.

(7) Tokic, M.; Hatzimanikatis, V.; Miskovic, L. Large-Scale Kinetic Metabolic Models of Pseudomonas Putida KT2440 for Consistent Design of Metabolic Engineering Strategies. Biotechnol. Biofuels 2020, 13, 33.

(8) Jouhten, P.; Konstantinidis, D.; Pereira, F.; Andrejev, S.; Grkovska, K.; Castillo, S.; Ghiachi, P.; Beltran, G.; Almaas, E.; Mas, A.; Warringer, J.; Gonzalez, R.; Morales, P.; Patil, K. R. Predictive Evolution of Metabolic Phenotypes Using Model-Designed Environments. Mol. Syst. Biol. 2022, 18 (10), e10980.

(9) Myers, P. J.; Lee, S. H.; Lazzara, M. J. MECHANISTIC AND DATA-DRIVEN MODELS OF CELL SIGNALING: TOOLS FOR FUNDAMENTAL DISCOVERY AND RATIONAL DESIGN OF THERAPY. Curr Opin Syst Biol 2021, 28. https://doi.org/10.1016/j.coisb.2021.05.010.

(10) Kim, O. D.; Rocha, M.; Maia, P. A Review of Dynamic Modeling Approaches and Their Application in Computational Strain Optimization for Metabolic Engineering. Front. Microbiol. 2018, 9, 1690.

(11) Benisch, M.; Benzinger, D.; Kumar, S.; Hu, H.; Khammash, M. Optogenetic Closed-Loop Feedback Control of the Unfolded Protein Response Optimizes Protein Production. Metab. Eng. 2023, 77, 32–40.

(12) Braniff, N.; Ingalls, B. New Opportunities for Optimal Design of Dynamic Experiments in Systems and Synthetic Biology. Current Opinion in Systems Biology 2018, 9, 42–48.

(13) Ilia, K.; Shakiba, N.; Bingham, T.; Jones, R. D.; Kaminski, M. M.; Aravera, E.; Bruno, S.; Palacios, S.; Weiss, R.; Collins, J. J.; Del Vecchio, D.; Schlaeger, T. M. Synthetic Genetic Circuits to Uncover and Enforce the OCT4 Trajectories of Successful Reprogramming of Human Fibroblasts. bioRxiv 2023. https://doi.org/10.1101/2023.01.25.525529.

(14) Kepler, T. B.; Elston, T. C. Stochasticity in Transcriptional Regulation: Origins, Consequences, and Mathematical Representations. Biophys. J. 2001, 81 (6), 3116–3136.

(15) Elowitz, M. B.; Levine, A. J.; Siggia, E. D.; Swain, P. S. Stochastic Gene Expression in a Single Cell. Science 2002, 297 (5584), 1183–1186.

(16) Cai, L.; Friedman, N.; Xie, X. S. Stochastic Protein Expression in Individual Cells at the Single Molecule Level. Nature 2006, 440 (7082), 358–362.

(17) Scott, M.; Klumpp, S.; Mateescu, E. M.; Hwa, T. Emergence of Robust Growth Laws from Optimal Regulation of Ribosome Synthesis. Mol. Syst. Biol. 2014, 10 (8), 747.

(18) Pedraza, J. M.; van Oudenaarden, A. Noise Propagation in Gene Networks. Science 2005, 307 (5717), 1965–1969.

(19) Gillespie, D. T. Exact Stochastic Simulation of Coupled Chemical Reactions. J. Phys. Chem. 1977, 81 (25), 2340–2361.

(20) Kolbe, N.; Hexemer, L.; Bammert, L.-M.; Loewer, A.; Lukáčová-Medvid’ová, M.; Legewie, S. Data-Based Stochastic Modeling Reveals Sources of Activity Bursts in Single-Cell TGF-β Signaling. PLOS Computational Biology. 2022, p e1010266. https://doi.org/10.1371/journal.pcbi.1010266.

(21) Loos, C.; Hasenauer, J. Mathematical Modeling of Variability in Intracellular Signaling. Current Opinion in Systems Biology 2019, 16, 17–24.

(22) Erdem, C.; Mutsuddy, A.; Bensman, E. M.; Dodd, W. B.; Saint-Antoine, M. M.; Bouhaddou, M.; Blake, R. C.; Gross, S. M.; Heiser, L. M.; Feltus, F. A.; Birtwistle, M. R. A Scalable, Open-Source Implementation of a Large-Scale Mechanistic Model for Single Cell Proliferation and Death Signaling. Nat. Commun. 2022, 13 (1), 3555.

(23) Covert, M. W.; Gillies, T. E.; Kudo, T.; Agmon, E. A Forecast for Large-Scale, Predictive Biology: Lessons from Meteorology. Cell Syst 2021, 12 (6), 488–496.

(24) Collombet, S.; van Oevelen, C.; Sardina Ortega, J. L.; Abou-Jaoudé, W.; Di Stefano, B.; Thomas-Chollier, M.; Graf, T.; Thieffry, D. Logical Modeling of Lymphoid and Myeloid Cell Specification and Transdifferentiation. Proc. Natl. Acad. Sci. U. S. A. 2017, 114 (23), 5792– 5799.

(25) Paulevé, L.; Kolčák, J.; Chatain, T.; Haar, S. Reconciling Qualitative, Abstract, and Scalable Modeling of Biological Networks. https://doi.org/10.1101/2020.03.22.998377.

(26) Le Novère, N. Quantitative and Logic Modelling of Molecular and Gene Networks. Nat. Rev. Genet. 2015, 16 (3), 146–158.

(27) Thomas, R. Boolean Formalization of Genetic Control Circuits. J. Theor. Biol. 1973, 42 (3), 563–585.

(28) Bintu, L.; Yong, J.; Antebi, Y. E.; McCue, K.; Kazuki, Y.; Uno, N.; Oshimura, M.; Elowitz, M. B. Dynamics of Epigenetic Regulation at the Single-Cell Level. Science 2016, 351 (6274), 720–724.

(29) Babtie, A. C.; Stumpf, M. P. H. How to Deal with Parameters for Whole-Cell Modelling. J. R. Soc. Interface 2017, 14 (133). https://doi.org/10.1098/rsif.2017.0237.

(30) Christodoulou, D.; Link, H.; Fuhrer, T.; Kochanowski, K.; Gerosa, L.; Sauer, U. Reserve Flux Capacity in the Pentose Phosphate Pathway Enables Escherichia Coli’s Rapid Response to Oxidative Stress. Cell Syst 2018, 6 (5), 569–578.e7.

(31) Kim, D. W.; Hong, H.; Kim, J. K. Systematic Inference Identifies a Major Source of Heterogeneity in Cell Signaling Dynamics: The Rate-Limiting Step Number. Sci Adv 2022, 8 (11), eabl4598.

(32) Vanlier, J.; Tiemann, C. A.; Hilbers, P. A. J.; van Riel, N. A. W. Parameter Uncertainty in Biochemical Models Described by Ordinary Differential Equations. Math. Biosci. 2013, 246 (2), 305–314.

(33) Persson, S.; Welkenhuysen, N.; Shashkova, S.; Wiqvist, S.; Reith, P.; Schmidt, G. W.; Picchini, U.; Cvijovic, M. Scalable and Flexible Inference Framework for Stochastic Dynamic Single-Cell Models. PLoS Comput. Biol. 2022, 18 (5), e1010082.

(34) Bar-Joseph, Z.; Gitter, A.; Simon, I. Studying and Modelling Dynamic Biological Processes Using Time-Series Gene Expression Data. Nat. Rev. Genet. 2012, 13 (8), 552–564.

(35) Chan, T. E.; Stumpf, M. P. H.; Babtie, A. C. Gene Regulatory Network Inference from Single-Cell Data Using Multivariate Information Measures. Cell Syst 2017, 5 (3), 251–267.e3.

(36) Sarmah, D.; Smith, G. R.; Bouhaddou, M.; Stern, A. D.; Erskine, J.; Birtwistle, M. R. Network Inference from Perturbation Time Course Data. NPJ Syst Biol Appl 2022, 8 (1), 42.

(37) Henriques, D.; Villaverde, A. F.; Rocha, M.; Saez-Rodriguez, J.; Banga, J. R. Data-Driven Reverse Engineering of Signaling Pathways Using Ensembles of Dynamic Models. PLoS Comput. Biol. 2017, 13 (2), e1005379.

(38) Treloar, N. J.; Braniff, N.; Ingalls, B.; Barnes, C. P. Deep Reinforcement Learning for Optimal Experimental Design in Biology. PLoS Comput. Biol. 2022, 18 (11), e1010695.

(39) Schmidt, M.; Lipson, H. Distilling Free-Form Natural Laws from Experimental Data. Science 2009, 324 (5923), 81–85.

(40) Beroza, G. C.; Segou, M.; Mostafa Mousavi, S. Machine Learning and Earthquake Forecasting-next Steps. Nat. Commun. 2021, 12 (1), 4761.

(41) Pilania, G.; Wang, C.; Jiang, X.; Rajasekaran, S.; Ramprasad, R. Accelerating Materials Property Predictions Using Machine Learning. Sci. Rep. 2013, 3, 2810.

(42) Pinheiro, G. A.; Mucelini, J.; Soares, M. D.; Prati, R. C.; Da Silva, J. L. F.; Quiles, M. G. Machine Learning Prediction of Nine Molecular Properties Based on the SMILES Representation of the QM9 Quantum-Chemistry Dataset. J. Phys. Chem. A 2020, 124 (47), 9854–9866.

(43) Ji, Y.; Lotfollahi, M.; Wolf, F. A.; Theis, F. J. Machine Learning for Perturbational Single-Cell Omics. Cell Syst 2021, 12 (6), 522–537.

(44) Ma, J.; Yu, M. K.; Fong, S.; Ono, K.; Sage, E.; Demchak, B.; Sharan, R.; Ideker, T. Using Deep Learning to Model the Hierarchical Structure and Function of a Cell. Nat. Methods 2018, 15 (4), 290–298.

(45) Baek, M.; Baker, D. Deep Learning and Protein Structure Modeling. Nat. Methods 2022, 19 (1), 13–14.

(46) Nikolados, E.-M.; Wongprommoon, A.; Aodha, O. M.; Cambray, G.; Oyarzún, D. A. Accuracy and Data Efficiency in Deep Learning Models of Protein Expression. Nat. Commun. 2022, 13 (1), 7755.

(47) Ling, H.; Samarasinghe, S.; Kulasiri, D. Novel Recurrent Neural Network for Modelling Biological Networks: Oscillatory p53 Interaction Dynamics. Biosystems. 2013, 114 (3), 191–205.

(48) Costello, Z.; Martin, H. G. A Machine Learning Approach to Predict Metabolic Pathway Dynamics from Time-Series Multiomics Data. NPJ Syst Biol Appl 2018, 4, 19.

(49) Yuan, B.; Shen, C.; Luna, A.; Korkut, A.; Marks, D. S.; Ingraham, J.; Sander, C. CellBox: Interpretable Machine Learning for Perturbation Biology with Application to the Design of Cancer Combination Therapy. Cell Syst 2021, 12 (2), 128–140.e4.

(50) Nilsson, A.; Peters, J. M.; Meimetis, N.; Bryson, B.; Lauffenburger, D. A. Artificial Neural Networks Enable Genome-Scale Simulations of Intracellular Signaling. Nat. Commun. 2022, 13 (1), 3069.

(51) Baranwal, M.; Clark, R. L.; Thompson, J.; Sun, Z.; Hero, A. O.; Venturelli, O. S. Recurrent Neural Networks Enable Design of Multifunctional Synthetic Human Gut Microbiome Dynamics. Elife 2022, 11. https://doi.org/10.7554/eLife.73870.

(52) Wang, S.; Fan, K.; Luo, N.; Cao, Y.; Wu, F.; Zhang, C.; Heller, K. A.; You, L. Massive Computational Acceleration by Using Neural Networks to Emulate Mechanism-Based Biological Models. Nat. Commun. 2019, 10 (1), 4354.

(53) Lugagne, J.-B.; Blassick, C. M.; Dunlop, M. J. Deep Model Predictive Control of Gene Expression in Thousands of Single Cells. bioRxiv, 2022, 2022.10.28.514305. https://doi.org/10.1101/2022.10.28.514305.

(54) Wang, P.; Robert, L.; Pelletier, J.; Dang, W. L.; Taddei, F.; Wright, A.; Jun, S. Robust Growth of Escherichia Coli. Curr. Biol. 2010, 20 (12), 1099–1103.

(55) Ong, N. T.; Tabor, J. J. A Miniaturized Escherichia Coli Green Light Sensor with High Dynamic Range. Chembiochem 2018, 19 (12), 1255–1258.

(56) Chait, R.; Ruess, J.; Bergmiller, T.; Tkačik, G.; Guet, C. C. Shaping Bacterial Population Behavior through Computer-Interfaced Control of Individual Cells. Nat. Commun. 2017, 8 (1), 1535.

(57) Endres, D. M.; Schindelin, J. E. A New Metric for Probability Distributions. IEEE Trans. Inf. Theory 2003, 49 (7), 1858–1860.

(58) Hartigan, J. A.; Hartigan, P. M. The Dip Test of Unimodality. Ann. Stat. 1985, 13 (1), 70–84.

(59) Lapin, A.; Perfahl, H.; Jain, H. V.; Reuss, M. Integrating a Dynamic Central Metabolism Model of Cancer Cells with a Hybrid 3D Multiscale Model for Vascular Hepatocellular Carcinoma Growth. Sci. Rep. 2022, 12 (1), 12373.

(60) Gupta, A.; Schwab, C.; Khammash, M. DeepCME: A Deep Learning Framework for Computing Solution Statistics of the Chemical Master Equation. PLoS Comput. Biol. 2021, 17 (12), e1009623.

(61) Sukys, A.; Öcal, K.; Grima, R. Approximating Solutions of the Chemical Master Equation Using Neural Networks. iScience 2022, 25 (9), 105010.

(62) Yang, J. H.; Wright, S. N.; Hamblin, M.; McCloskey, D.; Alcantar, M. A.; Schrübbers, L.; Lopatkin, A. J.; Satish, S.; Nili, A.; Palsson, B. O.; Walker, G. C.; Collins, J. J. A White-Box Machine Learning Approach for Revealing Antibiotic Mechanisms of Action. Cell 2019, 177 (6), 1649–1661.e9.

(63) Lundberg, S. M.; Nair, B.; Vavilala, M. S.; Horibe, M.; Eisses, M. J.; Adams, T.; Liston, D. E.; Low, D. K.-W.; Newman, S.-F.; Kim, J.; Lee, S.-I. Explainable Machine-Learning Predictions for the Prevention of Hypoxaemia during Surgery. Nat Biomed Eng 2018, 2 (10), 749–760.

(64) Tenachi, W.; Ibata, R.; Diakogiannis, F. I. Deep Symbolic Regression for Physics Guided by Units Constraints: Toward the Automated Discovery of Physical Laws. arXiv [astro-ph.IM], 2023. http://arxiv.org/abs/2303.03192.

(65) Mandracchia, B.; Hua, X.; Guo, C.; Son, J.; Urner, T.; Jia, S. Fast and Accurate sCMOS Noise Correction for Fluorescence Microscopy. Nat. Commun. 2020, 11 (1), 94.

(66) Kingma, D. P.; Ba, J. Adam: A Method for Stochastic Optimization. In ICLR 2015 - Conference Track Proceedings; 2015.

(67) Huber, P. J. Robust Estimation of a Location Parameter. Ann. Math. Stat. 1964, 35 (1), 73– 101.

