## Supporting Information for "Deep neural networks for predicting single cell responses and probability landscapes"

1. Supplementary Figures
2. Supplementary Methods
3. Supplementary References

### Supplementary Figures

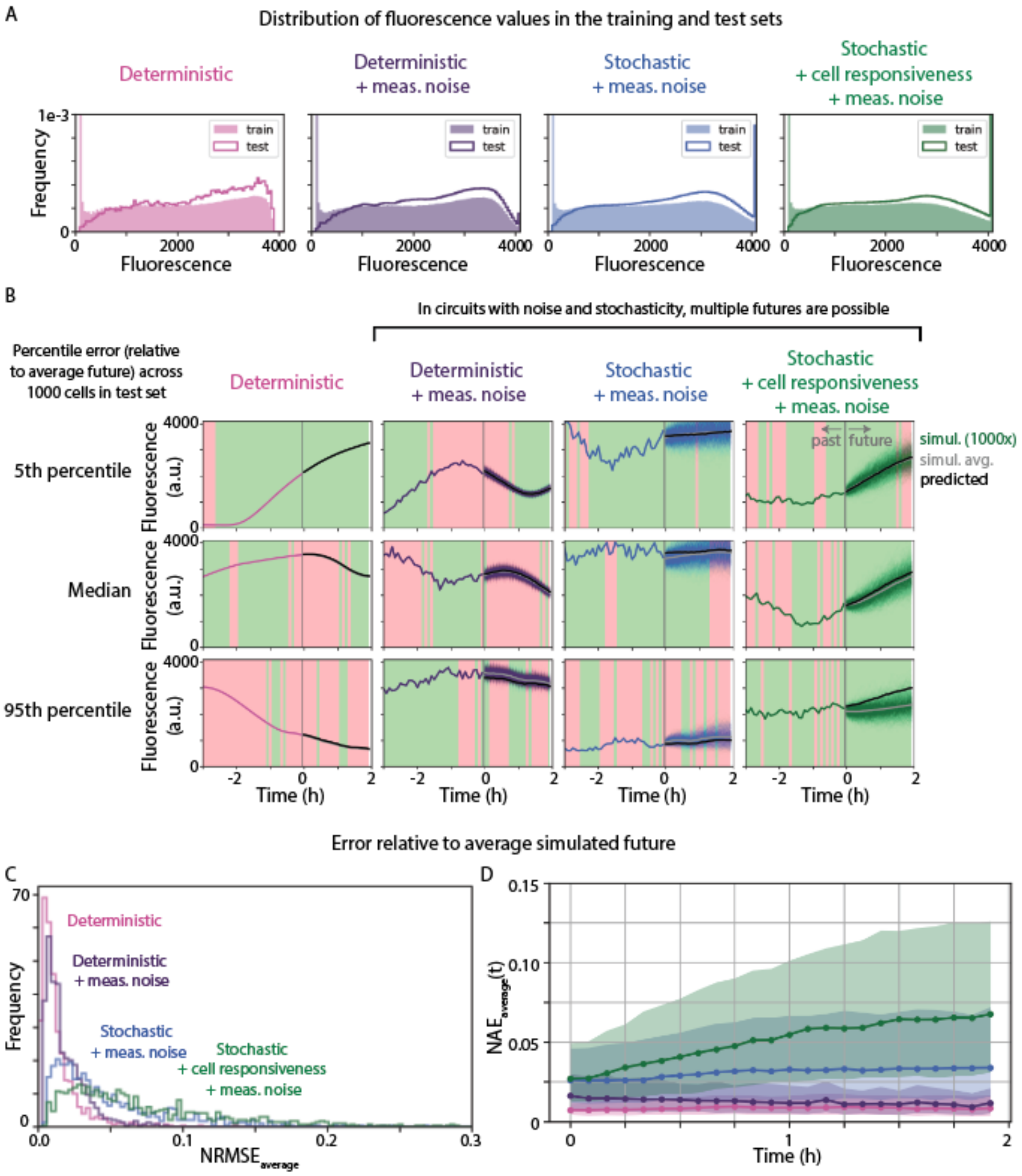


**Figure S1 - Prediction error varies across test set, relative to the spread of possible futures as well as the average future.**

(A) For both training and test data, cell responses are simulated in response to random sequences of red and green light. Training data sequences start with the cell in an "off" state, so the fluorescence distributions include more low reporter values. Otherwise, the distribution of fluorescence values in the two datasets are similar.

(B) The range of prediction error varies between the four models trained on different datasets. Each subplot shows a particular example from the test set, including the cell's past, 1000 simulated futures, average simulated futured (gray), and predicted future (black). The examples shown here represent different percentiles (5th, median, and 95th percentile) in one metric of prediction quality, specifically here the difference between the average simulated future (gray) and the predicted future (black).

(C) To quantify how well each model inferred average response dynamics, we computed the squared error between the average simulated future (gray in panel B) and the predicted future (black in panel B), normalized by the average response, and averaged over the prediction horizon ($NRMSE_{average}$, see Methods). Shown here are the distributions of error across the test set. Note that though models trained to predict stochastic responses have a larger spread of error, all four models have fairly similar and low median prediction errors.

(D) To show how prediction error evolved over time, we also computed the normalized absolute error ($NAE_{average}$, see Methods) between the average simulated future (gray in panel B) and the predicted future (black in panel B) at each point in time. Except for the model trained to predict responses with variation in cell responsiveness, all other models have fairly constant error, suggesting they correctly inferred the average dynamics of the cell response. Dots show the median value in the test set, and the shaded region spans the 25th to 75th percentile.

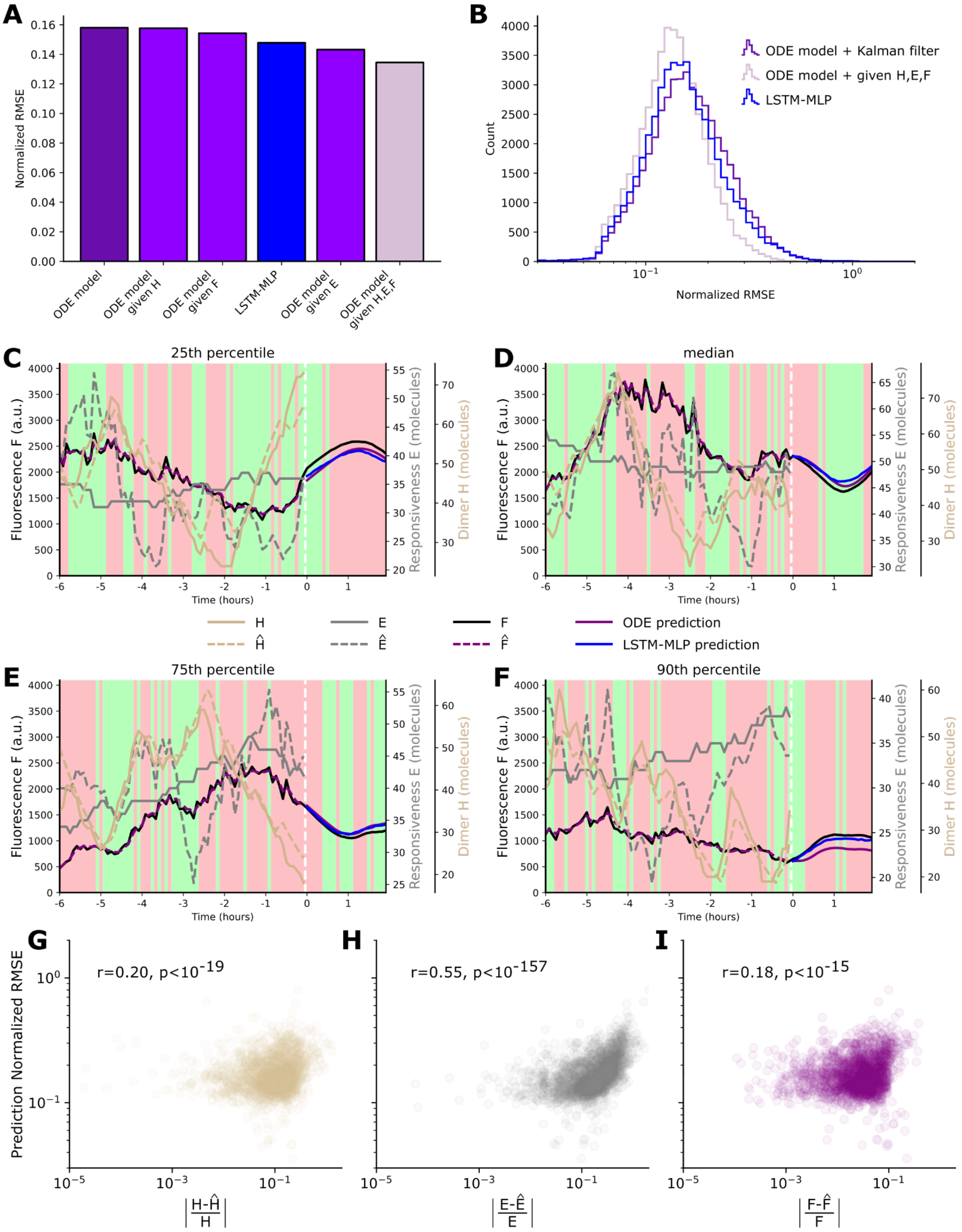


**Figure S2 - Comparison with state-of-the-art ODE model & hybrid Kalman filter.**

(A) Median of normalized RMSE for predictions from the LSTM-MLP and the ODE model with state estimation. When a variable *H*, *E*, or *F* is specified as given, its actual value was provided before prediction.

(B) Normalized RMSE distributions for LSTM-MLP, ODE model with Kalman filter, and ODE model with full state given.

(C) Prediction for the ODE model with Kalman filter at the 25th percentile of error. Before t = 0h, the black curve represents the measured fluorescence. Solid brown and gray curves represent actual counts for species H and E. The dashed purple curve represents "true" fluorescent protein counts estimated from the Kalman filter. Dashed brown and gray curves represent Kalman filter estimates for H and E. For t ,â• 0h, the solid black line represents the average of all 1000 simulated future fluorescence responses. The purple solid line represents the prediction from the ODE model based on estimated state. For reference, the blue line represents the prediction from the LSTM-MLP. We do not show the prediction from the ODE model given the full actual state because it overlaps almost perfectly with the black curve.

(D) Prediction for the ODE model with Kalman filter at median error.

(E) Prediction for the ODE model with Kalman filter at the 75th percentile of error.

(F) Prediction for the ODE model with Kalman filter at the 90th percentile of error.

(G) Normalized RMSE for ODE model prediction versus normalized absolute error between actual species count $H$ and estimated count $\hat{H}$ at t = 0h, and corresponding Spearman correlation coefficient r. Error in the estimation of the state variables correlates with prediction error, especially for species $E$, explaining the lower performance of the ODE model with Kalman filter state estimation (Supplementary Methods).

(H) Prediction normalized RMSE versus estimation error for species $E$ at t = 0h.

(I) Prediction normalized RMSE versus estimation error for species $F$ at t = 0h.

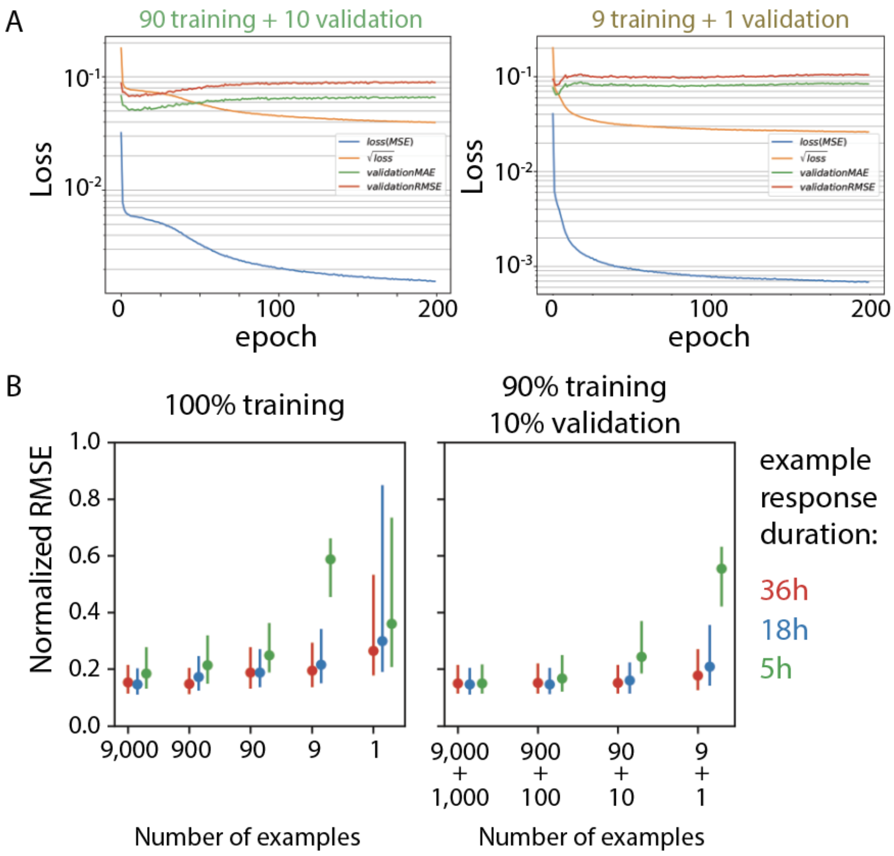


**Figure S3 - Validation set improves prediction quality for small training set sizes.**
Models were trained on different subsets of the data to explore how many and what types of examples can train a model that generalizes well to the test set.

(A) Training loss, quantified as mean-square error (MSE) and its square root, decreases with training epoch. By contrast, the validation error, shown as mean absolute error (MAE) and root mean square error (RMSE), at first decreases and then increases as the model overfits and loses generality. Training loss is shown for models trained on 90 training examples (10 validation examples) and 9 training examples (1 validation example).

(B) Models that minimize error in the training set (left plot) have lower prediction error (i.e. normalized RMSE) when more examples are used. Example responses that are longer duration also reduce prediction error. For the same number of examples, models that minimize error on the validation set (right plot) have the same or lower prediction error. Dots are median of normalized RSME in the test set, and the lines span the 25th to 75th percentiles.

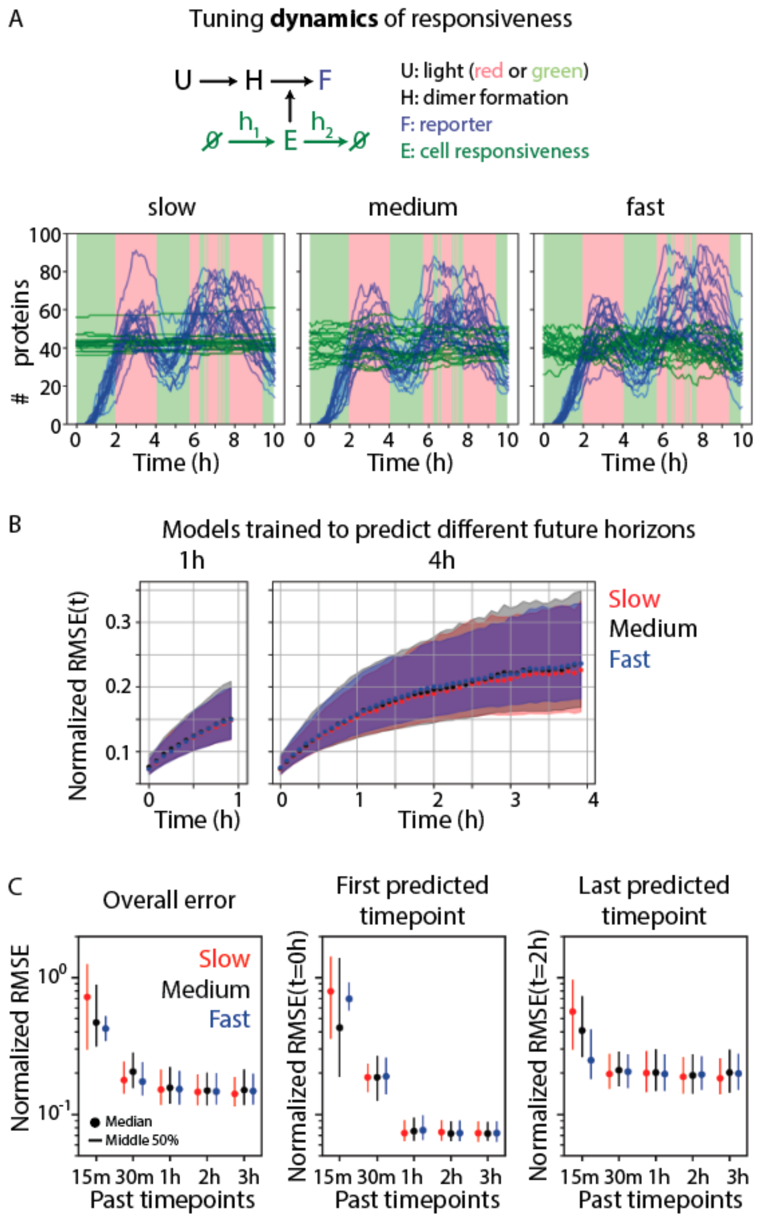


**Figure S4 - Increasing speed of fluctuations does not affect median prediction quality unless the amount of past information is very limited.**

(A) Sample stochastic simulations of cell responses to the same light input show the dynamics of reporter (*F*) and cell responsiveness (*E*) in the presence of different dynamics of *E*.

(B) Models were trained to predict either one or four hours into the future. Normalized RMSE(t) was computed across all examples in the test set; dots show the median, while the shaded regions span the 25th to 75th percentiles. Models trained on either slow, medium, or fast dynamics of *E* (see Supplementary Methods) have similar prediction error, across horizons or timepoints.

(C) Models were trained to predict the future with different numbers of past timepoints. No matter the dynamics of *E*, normalized RMSE decreased up until 1 hour of past response was provided. Endpoint prediction error is always higher than initial timepoint error, except when the model is trained on very few past timepoints. In each panel, dots show the median of the test set, while lines span the 25th to 75th percentiles.

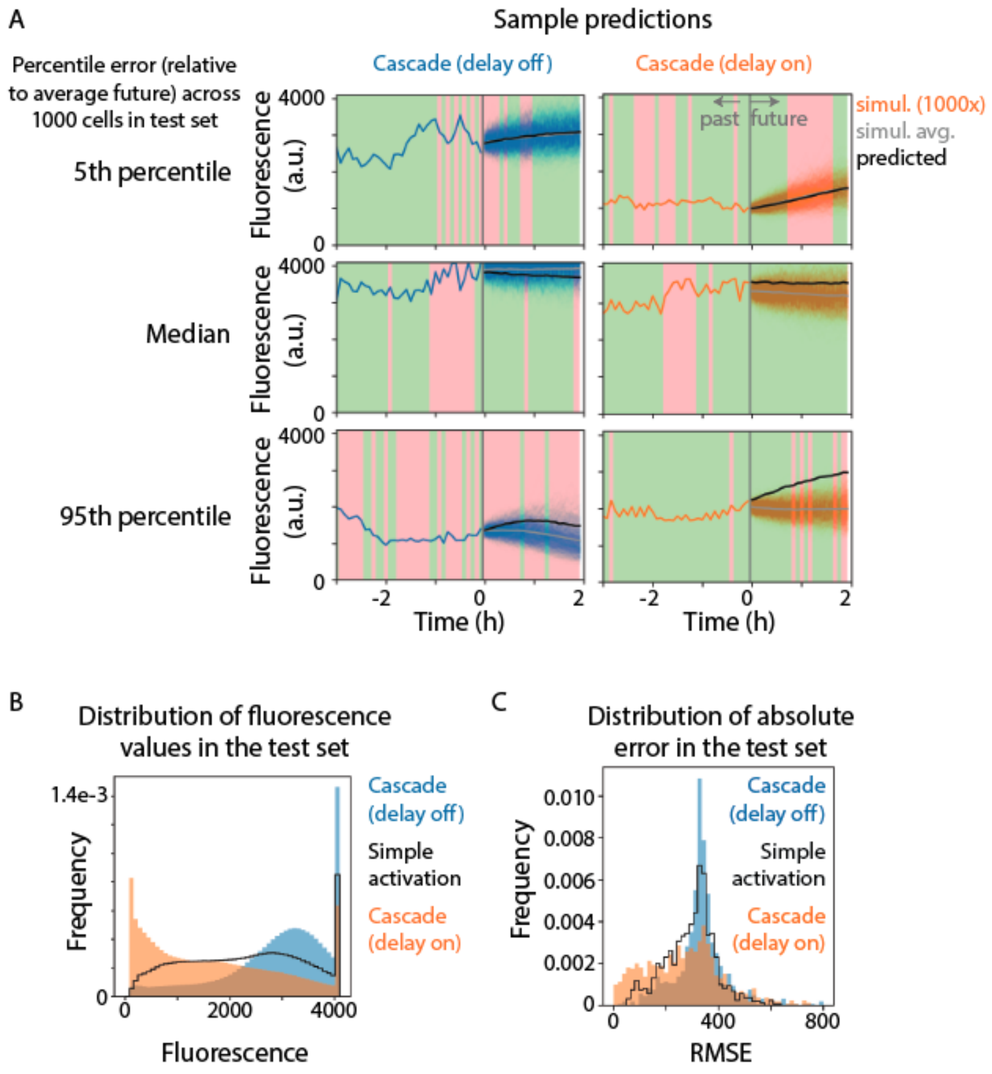


**Figure S5: LSTM-MLP can predict cascade response as well as simple activation response.**

(A) Models were trained to predict the dynamics of a cascade with slower off dynamics ("delay off", blue) and slower on dynamics ("delay on", orange). The 5th percentile, median, and 95th percentile predictions across the 1000 cells in the test set (based on squared error between the average simulated and predicted futures) are shown for both models. The vertical gray line at t = 0h separates the single simulated past from the 1000 simulated futures (blue or orange), average across those simulated futures (gray), and the model's predicted future (black).

(B) The distribution of fluorescence values in the test set shows that the simple activation circuit (black) evenly samples most fluorescence values, whereas the "delay off" and "delay on" cascades are biased towards higher and lower values respectively.

(C) The distribution of absolute error (i.e. RMSE) across the test set is similar for models trained on either the simple activation or cascade circuits. All circuits have similar maximum error, but the “delay on” cascade has lower absolute error, due to its lower responses in the test set.

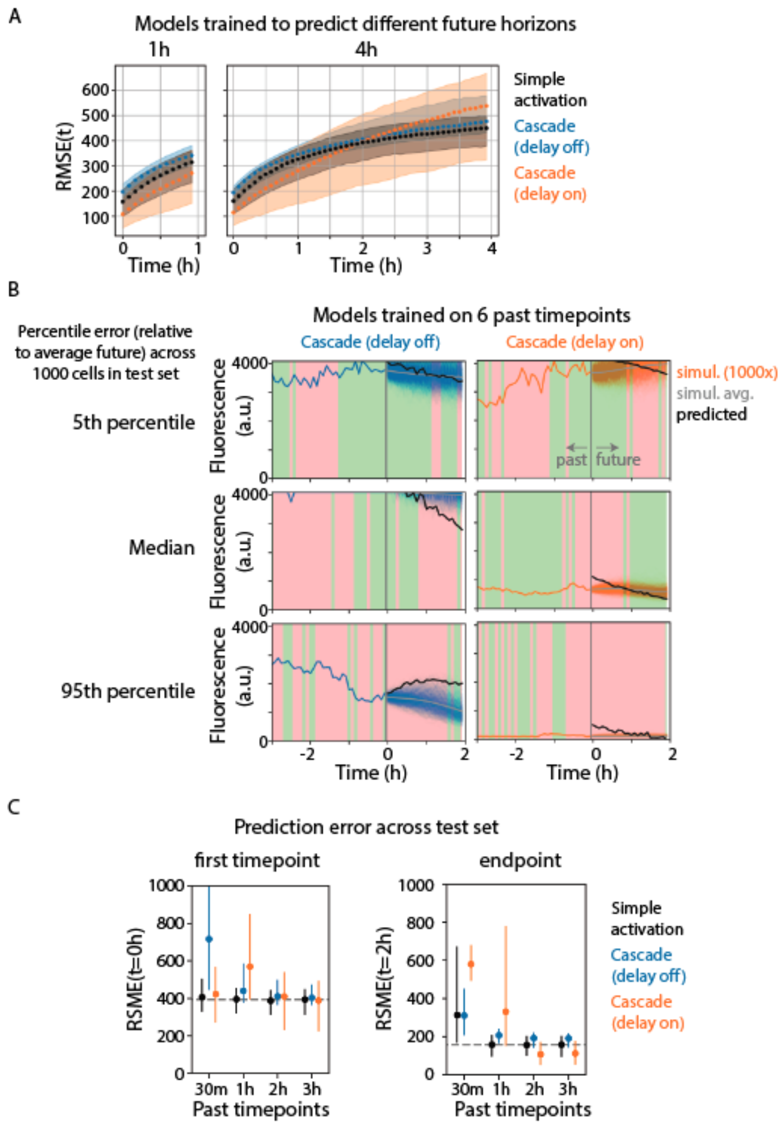


**Figure S6: Prediction error for cascade circuit is similar across future horizon, but more past information is required to predict cascade responses.**

(A) The delayed responses of the cascade circuit could make future dynamics easier to predict. We therefore compared the RMSE(t) of models trained to predict 1h or 4h into the future for the simple activation and the two cascade circuits. For less than an hour into the future, the absolute error tracks differences in the distribution of fluorescence values, with lower responses in the “delay on” cascade producing lower error. At longer timescales, the cascade circuits had higher and more broadly distributed errors.

(B) The cascade-associated delays can also increase the requirement for past information. Models trained to predict responses using only 30 minutes of past information rarely predicted responses well, as they could only accurately predict either the early or late response. Percentile responses taken from the distribution of error relative to the average future for each of the 1000 cells in the test set.

(C) To understand where prediction quality broke down when models had access to only a small part of the past response, we plotted the median (dot) and 25th to 75th percentiles (vertical line) of normalized RMSE values for models trained to predict future responses with 30 minutes, 1 hour, 2 hour, or 3 hours of past information. The dotted gray line represents the median of normalized RMSE for a model predicting simple activation responses using 3 hours of the past, the benchmark for models predicting other circuits and/or with less past information. Surprisingly, even though the initial timepoint should be the easiest to predict, models trained to predict the "delay on" cascade circuit had large errors at the initial timepoint.

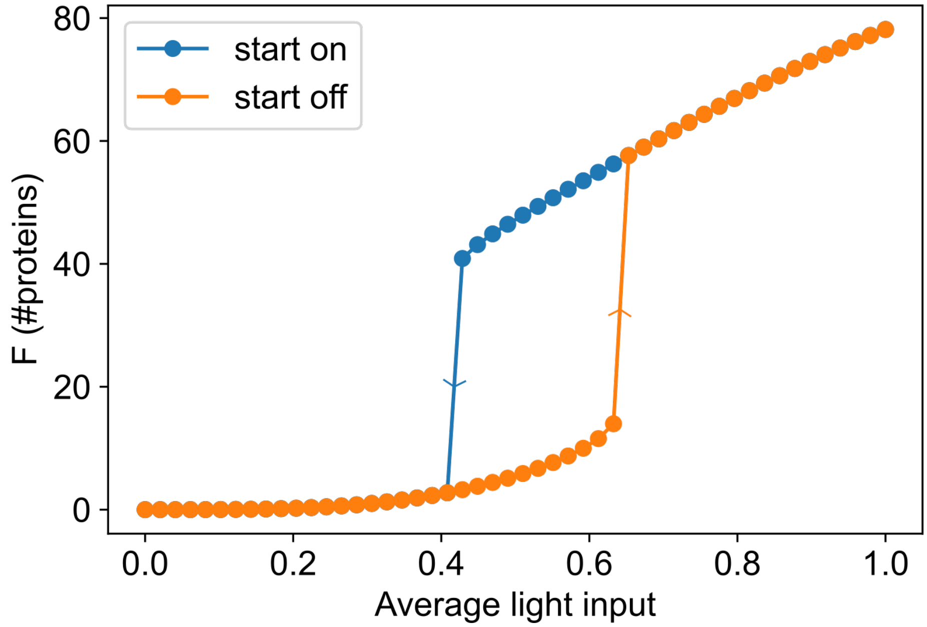


**Figure S7 - Hysteresis cycle of the auto-activation circuit.**
The deterministic solver was used to obtain the steady state response of the self-activating circuit to average light inputs ranging from 0 (fully red) to 1 (fully green). The steady state value was computed from two initial states where the system was either fully activated (blue dots) or fully repressed (orange dots). For intermediate light inputs, the system displays bistability.

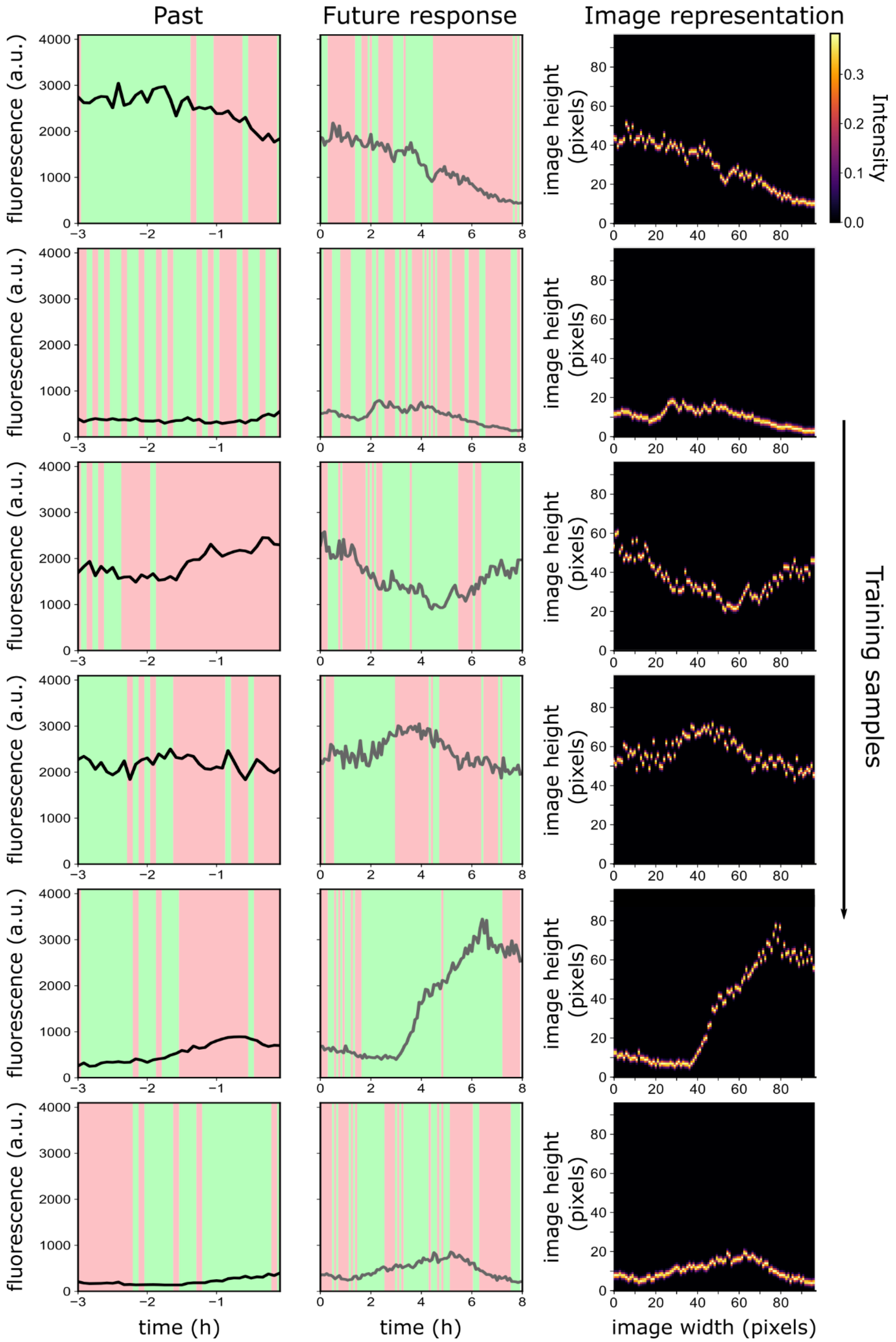


**Figure S8 - Training sample examples for the auto-activation circuit and illustration of the image representation used to train the convolutional decoder.**

Each row is a randomly-selected training sample from our dataset. The past of the cell is fed to the encoder, and the future cell response trajectory is formatted into an image-like representation of 96x96 pixels to use as ground truth against the prediction of the convolutional decoder.

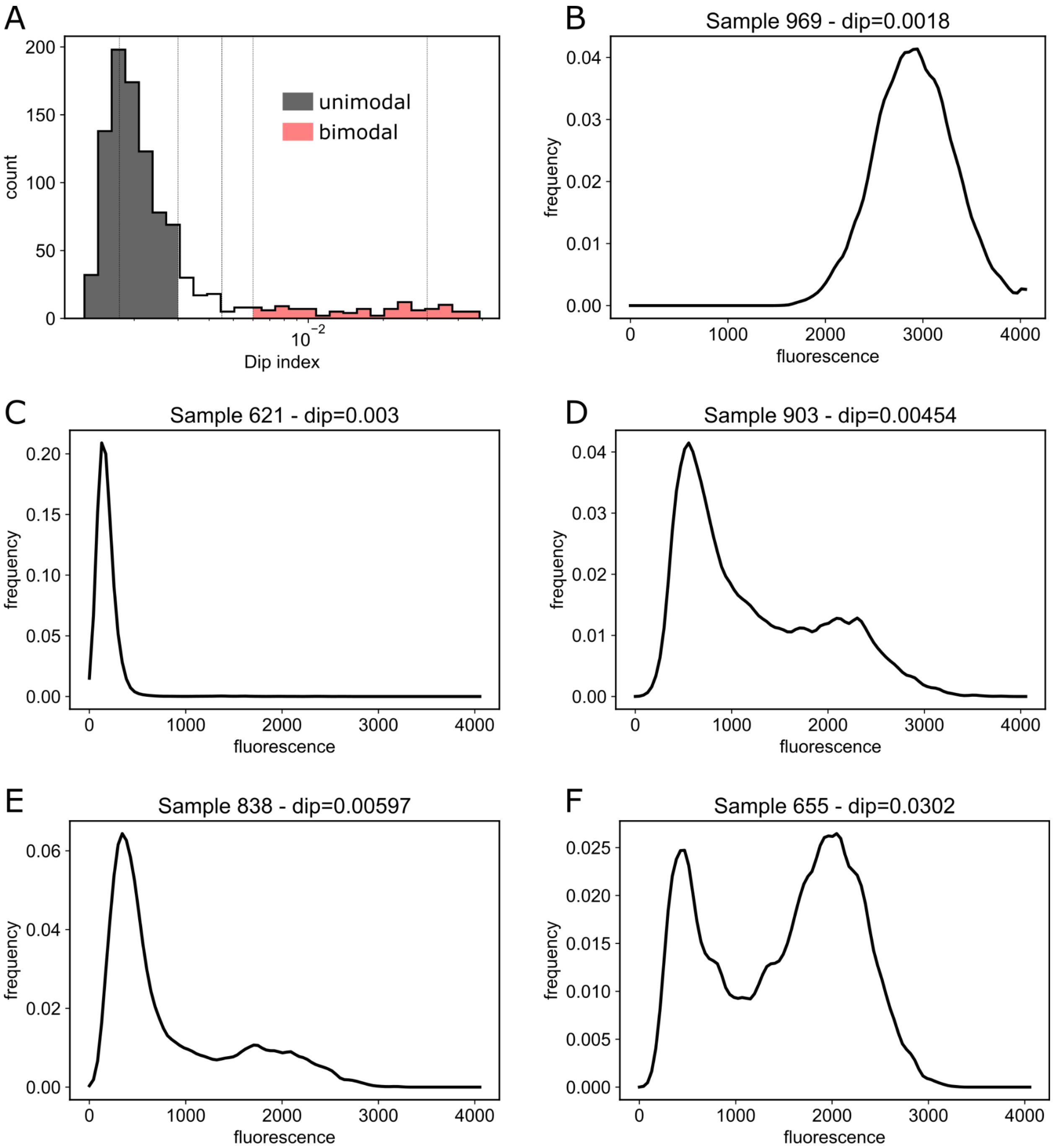


**Figure S9 - Results of the dip test results on our test set.**

(A) Histogram of dip index value over the test set. We considered samples under 0.003 to be unimodal, and samples above 0.006 to be bimodal. The vertical dashed lines represent the samples shown in the other panels.

(B-F) Fluorescence distributions over the last hour of the response for representative samples in the test set. Dip index value is listed in the title.

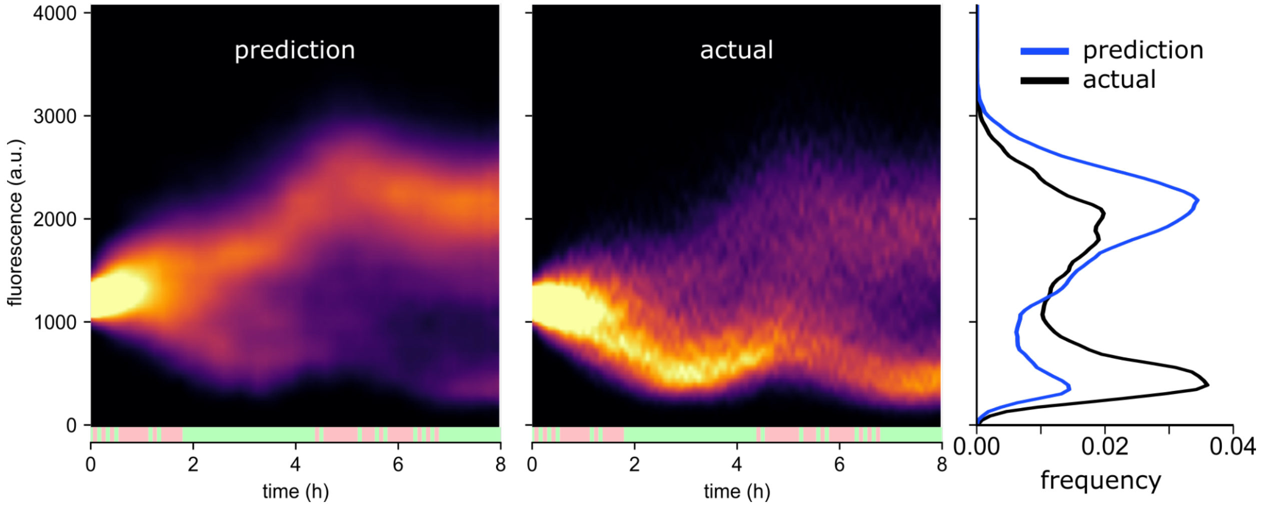


**Figure S10 - Worst prediction in our test set.**
An apparent error in state estimation leads the network to incorrectly predict which of the two modes will prevail throughout the cell response. The total Jensen-Shannon distance is 0.340.

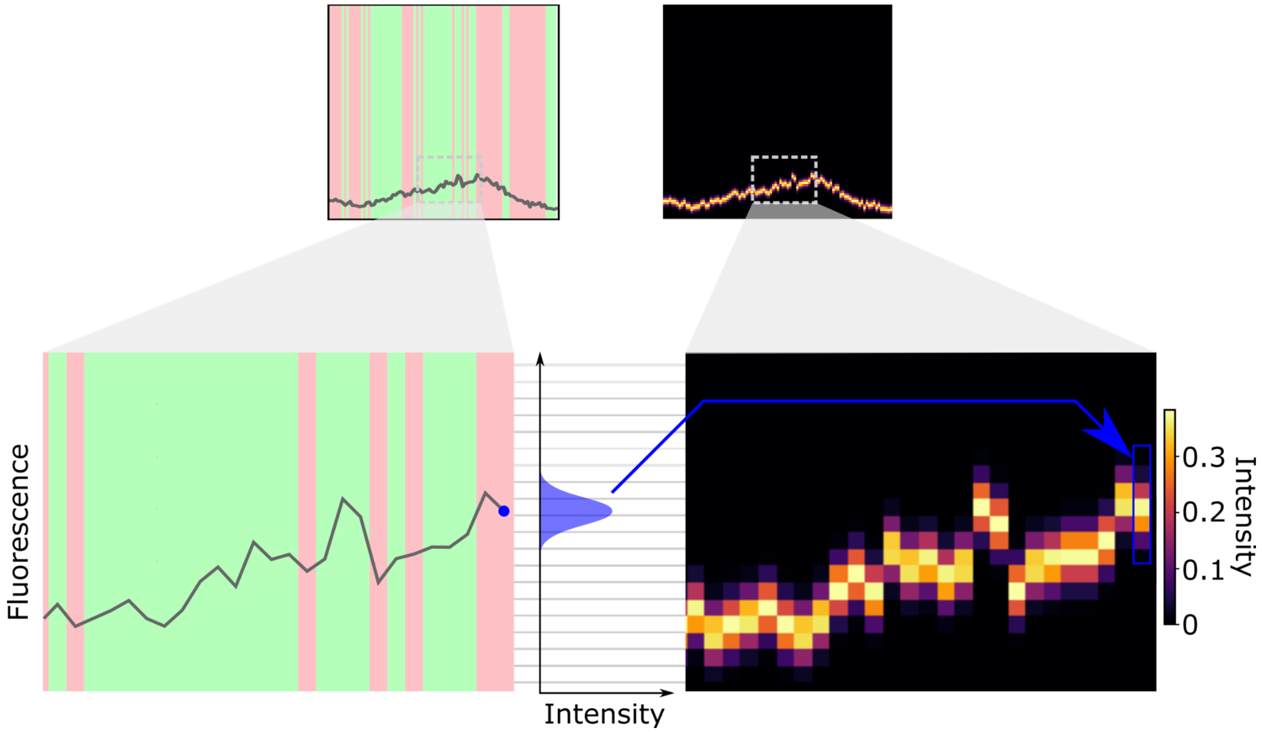


**Figure S11 - Gaussian kernel applied to fluorescence values to encode them into image pixels.**
Fluorescence values at each timepoint are convolved with a continuous Gaussian kernel. The resulting Gaussian curve is binned along the quantization thresholds (horizontal gray lines) and the area under the curve within each bin determines the pixel intensities for this timepoint. The cut-out shown in this figure is from the last sample shown in Figure S8.

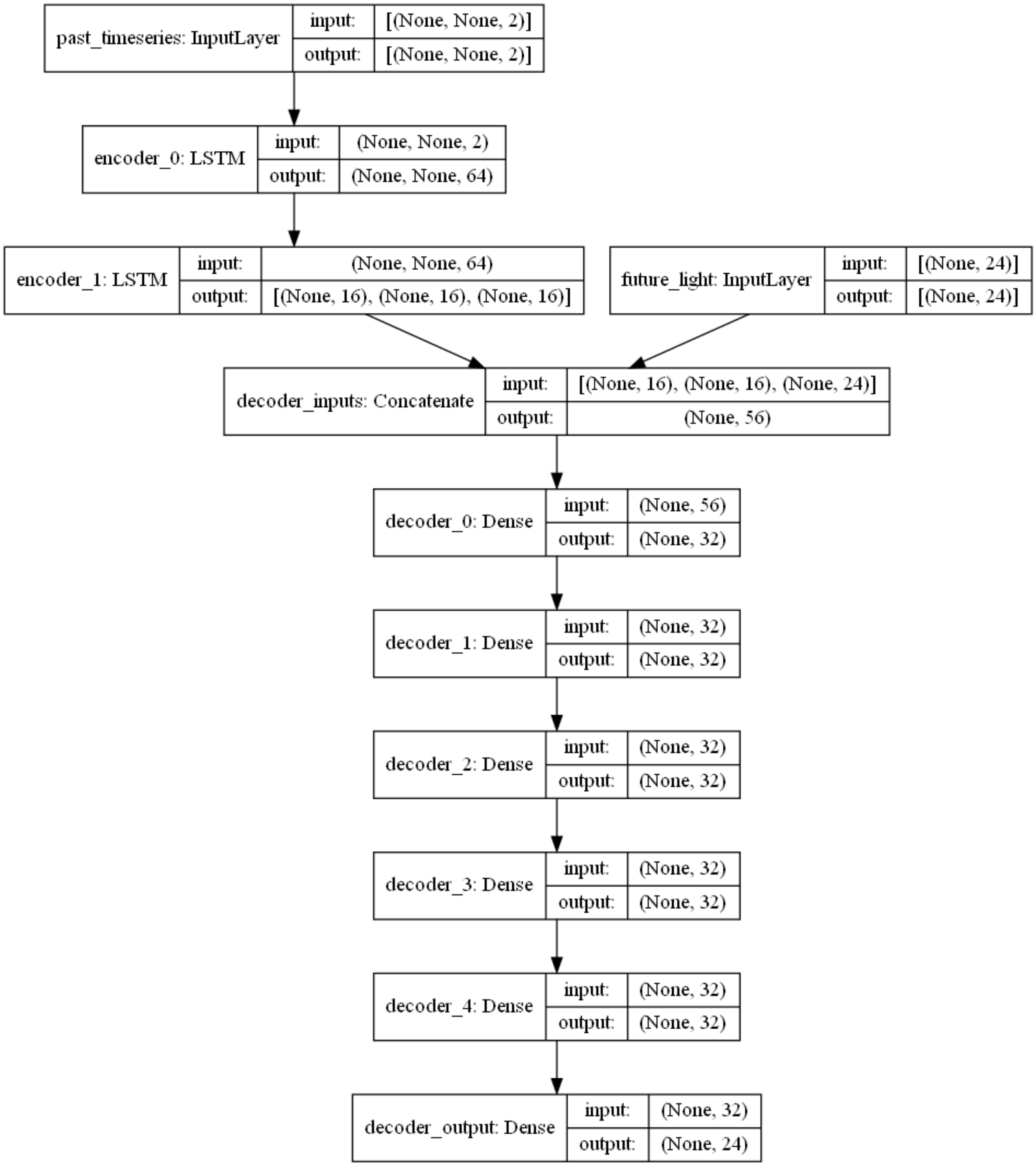


**Figure S12 - Architecture of the network with the MLP decoder.**
Layer class names refer to the Keras API names. The dimensions of the input and output tensors of each layer are shown on the right. The first dimension represents the batch size, and is set to None because batch size can be changed dynamically. For LSTM layers, the 2nd dimension is also None because the past time-series length can also vary. The past_timeseries InputLayer features two channels for the past fluorescence and past stimulations. Note that the hidden state and cell state of the last LSTM are passed to the decoder, and not the output tensor. This architecture is for the 2 hour prediction horizon, hence the 24-element future_light input tensor, and the 24-element decoder_output tensor.

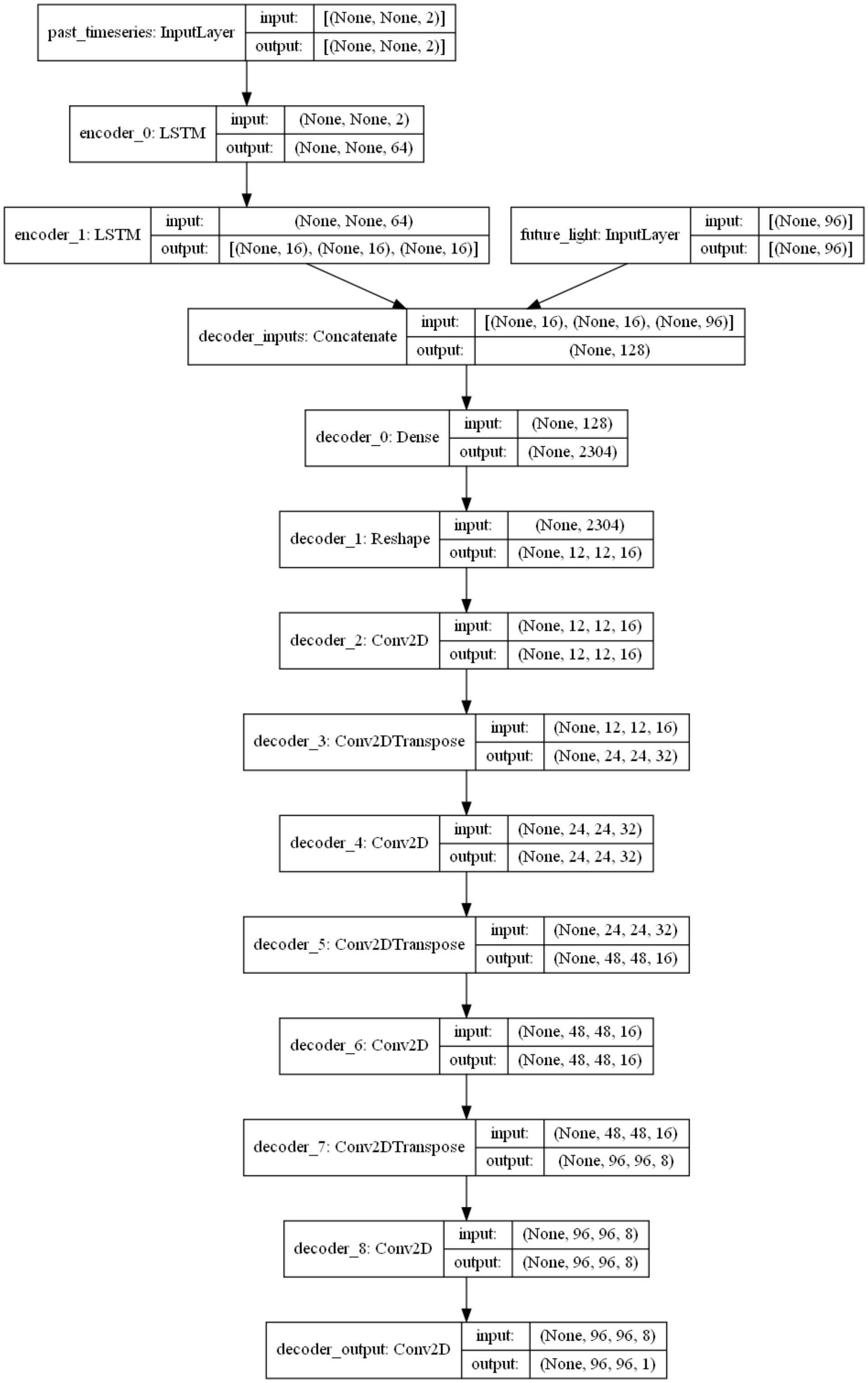


**Figure S13 - Architecture of the network with the convolutional decoder.**
Encoder architecture is the same for both the MLP and the convolutional decoder. For the convolutional decoder, the prediction horizon is 8 hours, leading to a future_light tensor of 96-element long vectors. The decoder_output tensor produces an image of shape 96x96, for the 96-timepoints horizon and the 96-binned quantization of fluorescence values.

### Supplementary Methods

#### Genetic circuit models

All circuit models can be simulated deterministically or stochastically. The reaction propensities are automatically re-formatted into an ODE set for deterministic simulations. All parameters are expressed in units of proteins, and time-related parameters are in (simulated) minutes or 1/minutes. Except where noted, parameters are taken from Chait et al.[1] All light inputs are applied with a time delay $\tau$ of 12 minutes between when the light is applied and the cell begins to respond (i.e. when species *U* is set). This approximates the dynamics of phosphorylation and dimerization that precede the formation of active CcaR dimer.

##### Simple circuit without cell responsiveness

This circuit simulates a fluorescent reporter *F* under the CcaSR optogenetic system. Cell-to-cell extrinsic responsiveness is not simulated here.

<u>Species:</u>
$\underline{U}$: Light input proxy species. This species is set by the optogenetic events with a delay of $\tau$.
$H$: CcaR dimer created in response to light inputs.
$F$: Reporter protein, under an activating Hill function of $H$.

<u>Reaction propensities:</u>

| | |
|---|---|
| ∅ → H | $\eta\ U$ |
| H → ∅ | $\nu\ H$ |
| ∅ → F | $\frac{a\ H^{n_h}}{K_H^{n_h} + H^{n_h}}$ |
| F → ∅ | $\nu\ F$ |

<u>Parameters:</u>

| $\eta$ | $\nu$ | $a$ | $K_H$ | $n_h$ | $\tau$ |
|---|---|---|---|---|---|
| 1/min | 0.01/min | 1.0/min | 45 | 3.6 | 12 min |

In the model by Chait et al.[1], the decay rate of the CcaR dimer ($c_2$) and of the fluorescent reporter (*b*) are different parameters. To simplify our model, we assumed that all proteins have the same decay rate, which is set by the dilution associated with cell division every 30 minutes. This yields a dilution rate of 0.01, which is very close to the reported best fit value for *b*, 0.0104. Also for simplicity, we increased *a*, the maximum production rate of *F*, to 1 from the published best fit value 0.2827. Lastly, the published fit model has a different expression for the Hill activation of *F*, such that our parameter $K_H$ is equivalent to their fit $\frac{\kappa}{c_2} \approx 7.8$. Since this parameter sets the sensitivity of the system to numbers of molecules of *H,* we chose a larger value for the default circuit model, to allow us to explore higher and lower sensitivity in other circuits.

#### Simple circuit with cell responsiveness

This circuit simulates a fluorescent reporter *F* under the CcaSR optogenetic system, but adds a species *E* that simulates cell-to-cell extrinsic responsiveness.

Species:
$E$: Extrinsic responsiveness proxy molecule that influences protein expression.
$U$: Light input proxy species. This species is set by the optogenetic events with a delay of $\tau$.
$H$: CcaR dimer created in response to light inputs.
$F$: Reporter protein, under an activating Hill function of $H$.

Reaction propensities:

| | |
|---|---|
| ∅ → E | $h_1$ |
| E → ∅ | $h_2\ E$ |
| ∅ → H | $\eta\ U$ |
| H → ∅ | $\nu\ H$ |
| ∅ → F | $\frac{a\ E\ H^{n_h}}{K_H^{n_h} + H^{n_h}}$ |
| F → ∅ | $\nu\ F$ |

Parameters:

| $\eta$ | $\nu$ | $h_1$ | $h_2$ | $a$ | $K_H$ | $n_h$ | $\tau$ |
|---|---|---|---|---|---|---|---|
| 1/min | 0.01/min | 0.04/min | 0.001/min | 0.025/min | 45 | 3.6 | 12 min |

These parameters are the "medium" dynamics of *E* shown in Figure S4. For the "slow" dynamics, $h_1$= 0.008 and $h_2$ = 0.0002. For the "fast" dynamics, $h_1$= 0.2 and $h_2$ = 0.005. We used a value for $h_2$ an order of magnitude smaller than the published best fit parameters[1] because these slow changes in responsiveness better matched what we had previously observed in our CcaSR experiments.[2]

#### Cascade

This circuit adds an intermediate activator species *I*. This intermediate is under the CcaSR optogenetic system, and then activates the reporter species *F*.

Species:
$E$: Extrinsic responsiveness proxy molecule that influences protein expression.
$U$: Light input proxy species. This species is set by the optogenetic events with a delay of $\tau$.
$H$: CcaR dimer created in response to light inputs.
$I$: Intermediate protein, under an activating Hill function of $H$.

$F$: Reporter protein, under an activating Hill function of $I$.

Reaction propensities:

| | |
|---|---|
| ∅ → E | $h_1$ |
| E → ∅ | $h_2\ E$ |
| ∅ → H | $\eta\ U$ |
| H → ∅ | $\nu\ H$ |
| ∅ → I | $\frac{a_I\ a\ E\ H^{n_h}}{K_H^{n_h} + H^{n_h}}$ |
| I → ∅ | $\nu\ I$ |
| ∅ → F | $\frac{a_F\ a\ E\ I^{n_i}}{K_I^{n_i} + I^{n_i}}$ |
| F → ∅ | $\nu\ F$ |

Parameters:

| $\eta$ | $\nu$ | $h_1$ | $h_2$ | $a$ | $K_H$ | $n_h$ | $K_I$ | $n_i$ | $a_I$ | $a_F$ | $\tau$ |
|---|---|---|---|---|---|---|---|---|---|---|---|
| 1/min | 0.01/min | 0.04/min | 0.001/min | 0.025/min | 45 | 3.6 | 70 | 3.6 | 1/min | 1.1204/min | 12 min |

These values are for the "delay on" case. For the "delay off" case, $K_I = 20$ and $a_F = 0.8408$. By changing $a_F$ alongside $K_I$, we maintain the same steady-state level of $F$ when $U$=1.

#### Autoactivation

This circuit is derived from the simple circuit without extrinsic cell responsiveness, and adds self-activation by the reporter species *F*. Cell-to-cell extrinsic responsiveness is not simulated here. Parameters $K_H$ and $K_F$ were selected such that the system displayed hysteresis for intermediate light inputs.

Species:
$U$: Light input proxy species. This species is set by the optogenetic events with a delay of $\tau$.
$H$: CcaR dimer created in response to light inputs.
$F$: Reporter protein, under an activating Hill function of $H$ and a self-activating Hill function.

Reaction propensities:

| | |
|---|---|
| ∅ → H | $\eta\ U$ |
| H → ∅ | $\nu\ H$ |
| ∅ → F | $\frac{a/2\ H^{n_h}}{K_H^{n_h} + H^{n_h}}$ |

| | |
|---|---|
| ∅ → F | $\frac{a/2\ F^{n_f}}{K_F^{n_f} + F^{n_f}}$ |
| F → ∅ | $\nu\ F$ |

<u>Parameters:</u>

| $\eta$ | $\nu$ | $K_H$ | $a$ | $n_h$ | $n_f$ | $K_F$ | $\tau$ |
|---|---|---|---|---|---|---|---|
| 1/min | 0.01/min | 90 | 1.0/min | 3.6 | 3.6 | 30 | 12 min |

#### ODE model and state estimation

Our stochastic simulations of the simple circuit with cell responsiveness are based on the ODE model described by Chait et al.[1] to forecast the fluorescence response of the CcaSR system in *E. coli*. In their study, they used this model to implement real-time control of gene expression in a model predictive framework. In order to be able to predict gene expression they first needed to estimate the state of the system, i.e. estimate the values of not only "true" fluorescence once noise is filtered out, but also of the hidden variables for the number of molecules of the dimer *H* and cell responsiveness *E*. To do this, they used moment closure on the chemical master equation to establish a set of differential equations[3] that model not only the evolution of the first order moments of the random variables for the species counts, i.e. their expected values, but also of their second order moments, so that they can predict the evolution of the covariance matrix of the system. This means that they can use a hybrid Kalman filter approach to estimate the state of the system at the present timepoint, and then use these estimated state values to predict the evolution of the system at future timepoints. The ODE set below is adapted from their study, with Equations 1-3 directly recognizable from the reaction propensities for the system, and Equations 4-6 for second order moments dynamics derived via moment closure:

$$\frac{\partial}{\partial t}H(t) = u(t-\tau) - \nu \cdot H(t) \tag{1}$$

$$\frac{\partial}{\partial t}\mathbb{E}[E(t)] = h_1 - h_2 \cdot \mathbb{E}[E(t)] \tag{2}$$

$$\frac{\partial}{\partial t}\mathbb{E}[F(t)] = a \cdot L(t) \cdot \mathbb{E}[E(t)] - \nu \cdot \mathbb{E}[F(t)] \tag{3}$$

$$\frac{\partial}{\partial t}\mathbb{E}[E(t)^2] = h_1 + 2h_1 \cdot \mathbb{E}[E(t)] + h_2 \cdot \mathbb{E}[E(t)] - 2h_2 \cdot \mathbb{E}[E(t)^2] \tag{4}$$

$$\frac{\partial}{\partial t}\mathbb{E}[E(t)F(t)] = h_1 \cdot \mathbb{E}[F(t)] + a \cdot L(t) \cdot \mathbb{E}[E(t)^2] - (h_2 + \nu) \cdot \mathbb{E}[E(t)F(t)] \tag{5}$$

$$\frac{\partial}{\partial t}\mathbb{E}[F(t)^2] = \nu \cdot \mathbb{E}[F(t)] + a \cdot L(t) \cdot \mathbb{E}[E(t)] + 2a \cdot L(t) \cdot \mathbb{E}[E(t)F(t)] - 2\nu \cdot \mathbb{E}[F(t)^2] \tag{6}$$

with $L(t) = \frac{H(t)^{n_H}}{{K_H}^{n_H} + H(t)^{n_H}}$

Note that *H(t)*, which represents the activation dynamics of the CcaSR system and incorporates a time delay on the optogenetic stimulations *u(t)*, is treated as fully deterministic: This is because the non-linearity of $L(t)$ makes moment closure analytically intractable otherwise. This is the only "erroneous" assumption in this model compared to our implementation of the stochastic simulations, where the reactions for the creation and deletion of species $H$ are stochastically simulated. $E(t)$ is a stochastic variable that represents the cell-to-cell variability in the "responsiveness" of the system. $F(t)$ is the number of fluorescent molecules and is also a stochastic variable. We converted between the range of numbers of fluorescent molecules and the range of measured fluorescence (multiplication factor 40 a.u./molecule and additive offset of 100 a.u.) as necessary for state estimation or to evaluate prediction accuracy (See Methods section in Main Text).

With this set of ODEs, future responses can be predicted, but also the state of the system can be estimated from past data with a hybrid Kalman filter. Briefly, the state estimator uses the model of the system to separate technical measurement noise from fluorescence fluctuations caused by gene expression variability, and estimates the true fluorescent protein counts ($F(t)$) as well as hidden variables for cell responsiveness ($E(t)$) and CcaSR activation ($H(t)$) levels at each time point by comparing model predictions to measurements. Because we are using a Kalman filter approach, technical noise is assumed to be additive, time-invariant Gaussian noise with mean zero and variance $R$. Because the prior for the covariance matrix of the system can be calculated with Equations 4-6, we can estimate how much of the fluctuations in measured fluorescence levels can be attributed to noise or to actual changes in the system's variables. By tuning the value for $R$, we can adjust the amount of noise rejection from the filter, i.e. how much the filter should "trust" the model vs. measurements. We evaluated a broad range of values and found that $R$=10 produced the most accurate predictions. We refer the reader to Chait et al.[1] for a detailed mathematical description of the state estimator algorithm, and to our code repository for a Python implementation of that algorithm.

Interestingly, even though this ODE model is based on almost perfect knowledge of the simulated system, it did not outperform our neural network in prediction accuracy (Figure S2A-B). Since our data is entirely simulated, we were able to investigate this surprising result. We found that, while in most cases state estimation was fairly accurate, the worst ODE predictions tended to arise from errors in state estimation (Figure S2C-F). Indeed, after we provided the actual state values to replace estimated values, ODE model predictions improved (Figure S2A-B). The ODE-based model became more accurate than the LSTM-MLP model only once given the actual count for species $E$ or the full state. We note that when given the full state, the prediction from the ODE model matches almost perfectly the average of all 1000 future responses, but the median of the normalized prediction RMSE across realizations is 0.134 because of the stochasticity of single responses. Finally, we show that there is a correlation between state estimation error and prediction error (Figure S2G-I), especially for species $E$. Finally, we note that computation time for ODE-based predictions is 3 to 4 orders of magnitude slower than for our LSTM-MLP model.

These results can be surprising given that the ODE model is based on perfect knowledge of the system dynamics and parameters. However, this is a difficult estimation problem, as the process is fundamentally stochastic, and species $E$ is independent from the rest of the system. Moreover, Kalman filtering relies on the assumption that system and measurement noise follow a Gaussian distribution, which is not the case in our simulated data and more generally when dealing with gene expression data. Similar Bayesian filtering approaches such as particle filters do not make this assumption and could perform better, however they rely on Monte Carlo simulations and are typically much slower than ODE and Kalman filter-based approaches.

**Supplementary References**


(1) Chait, R.; Ruess, J.; Bergmiller, T.; Tkačik, G.; Guet, C. C. Shaping Bacterial Population Behavior through Computer-Interfaced Control of Individual Cells. *Nat. Commun.* **2017**, *8* (1), 1535.
(2) Lugagne, J.-B.; Blassick, C. M.; Dunlop, M. J. Deep Model Predictive Control of Gene Expression in Thousands of Single Cells. *bioRxiv*, 2022, 2022.10.28.514305. https://doi.org/10.1101/2022.10.28.514305.
(3) Engblom, S. Computing the Moments of High Dimensional Solutions of the Master Equation. *Appl. Math. Comput.* **2006**, *180* (2), 498–515.
